# Therapy resistance in AML is mediated by cytoplasmic sequestration of the transcriptional repressor IRF2BP2

**DOI:** 10.1101/2025.05.22.655396

**Authors:** Mark J Althoff, Mohd Minhajuddin, Brett M. Stevens, Austin E. Gillen, Stephanie Gipson, Sweta B. Patel, Ian T. Shelton, Regan Miller, Ana Vujovic, Anna Krug, Tracy Young, William M. Showers, Monika Dzieciatkowska, Daniel Stephenson, Anit Tyagi, Jana M. Ellegast, Tristen Wright, Kimberly Stegmaier, Joseph T Opferman, Angelo D’Alessandro, Clayton A Smith, Craig T Jordan

**Affiliations:** Division of Hematology, University of Colorado Anschutz Medical Campus, Aurora, Colorado; RefinedScience, Aurora, Colorado; Department of Biochemistry and Molecular Genetics, University of Colorado Anschutz Medical Campus, Aurora, Colorado; Department of Cell and Molecular Biology, St. Jude Children’s Research Hospital, Memphis, Tennessee; Department of Pediatric Oncology, Dana-Farber Cancer Institute, Boston, Massachusetts; Division of Hematology/Oncology, Boston Children’s Hospital, Boston, Massachusetts; Broad Institute of MIT and Harvard, Cambridge, Massachusetts; Faculty of Medicine, University of Zurich, Zurich, Switzerland; Department of Medical Oncology and Hematology, University Hospital Zurich, Zurich, Switzerland

**Keywords:** Acute Myeloid Leukemia, fatty acid metabolism, therapy resistance, MCL1, IRF2BP2

## Abstract

While the development of venetoclax with azacitidine (ven/aza) has improved AML therapy, drug resistance remains a major challenge. Notably, primary ven/aza-resistant AML are frequently reliant on MCL1, however, the underlying mechanisms remain unclear. Co-immunoprecipitation of MCL1 from ven/aza-resistant AML samples coupled with mass spectrometry analysis identified the transcriptional repressor Interferon Regulatory Factor 2 Binding Protein 2 (IRF2BP2) as an MCL1 binding partner. This interaction results in cytoplasmic IRF2BP2 localization and loss of transcriptional repression within ven/aza-resistant leukemic stem cells (LSC). Consequently, ven/aza-resistant LSC have increased IRF2BP2 target gene expression, including acyl-CoA synthetase long-chain family member 1 (*ACSL1*), an essential rate-limiting enzyme for fatty acid oxidation (FAO). Inhibition of ACSL1 functionally impaired ven/aza-resistant LSC through a depletion of long-chain acyl-carnitine metabolites and FAO. Collectively, these data provide evidence for a previously undescribed mechanism by which MCL1 mediates IRF2BP2 cytoplasmic sequestration and consequent de-repression of *ACSL1*, thereby promoting ven/aza-resistance in AML.

## Introduction

Despite extensive efforts to develop improved therapies for acute myeloid leukemia (AML), patients often exhibit poor clinical outcomes. In particular, the development of clinical strategies to eradicate disease-initiating leukemia stem cells (LSC), continues to be a major challenge^1^. Early studies characterized LSCs as a seemingly homogeneous entity, but numerous subsequent reports have demonstrated that LSCs can be highly heterogeneous^2-4^. Perhaps most importantly, varying responsiveness to therapeutic challenge is clearly evident amongst LSCs of differing genotype and phenotype^2,5,6^. Thus, we and others have sought to better understand the central mechanisms that drive clinical response at the LSC level. These efforts have shown that aberrant metabolic properties are a central feature of primary human LSC populations and their response to therapy^6-12^. Specifically, the mechanism driving oxidative phosphorylation (OXPHOS) is fundamentally changed in LSCs, where the catabolism of amino acids is the primary driver of OXPHOS^8^, in contrast to normal hematopoietic stem cells (HSC) which mainly utilize glucose to support OXPHOS. Furthermore, LSCs are largely unable to compensate for inhibition of OXPHOS^7,13,14^, unlike most cell types which can employ glycolysis as an alternative means to produce ATP. Thus, strategies that target LSC-specific mechanisms of driving OXPHOS are an appealing therapeutic strategy.

To date, the best characterized clinical modulator of OXPHOS in AML is the BCL-2 inhibitor venetoclax (ven). As we previously demonstrated, 60-70% of newly diagnosed AML patients are sensitive to ven-mediated inhibition of OXPHOS in the LSC population^9,15-17^. The central mechanism of suppression involves reduction of amino acid catabolism^8^. Unfortunately, most patients who initially respond to ven-based therapy will relapse. Notably, although ven-based regimens directly target LSCs^8,9^, we have recently discovered that ven-resistant LSC, that either co-reside or evolve from ven-sensitive LSC^2,5^ confer resistance and disease progression. Fatty acid oxidation (FAO) has been implicated as an essential metabolic pathway in AML and is also evident in ven-resistant LSCs^6,18-21^. We have reported that activation of FAO can circumvent the effects of venetoclax and permit OXPHOS to proceed^6,8^. Consequently, understanding the mechanisms that activate FAO and developing strategies to suppress FAO in a leukemia-specific fashion have become a key objective in the field.

An additional component of venetoclax resistance is the relative activity of BCL2. As expected, BCL2 expression is relatively high in ven-sensitive AML cells and is reduced in resistance. Notably, several reports have described increased expression of MCL1, a related member of the BCL2 family, in AML patients where BCL2 is down-regulated^5,22-25^. In addition, we have reported that MCL1 appears to supplant the role of BCL2 in supporting FAO activity and OXPHOS in ven-resistant LSCs^5,25^. Thus, in the present study we investigated the molecular mechanisms by which MCL1 controls the metabolic phenotype found in ven-resistant LSCs. Our studies indicate that a previously undescribed role for MCL1 in sequestering the transcriptional repressor IRF2BP2 is central to mediating response to venetoclax in AML.

## Results

### MCL1 expression is associated with ven/aza-resistance

We and others have previously reported a distinct subset of ven/aza-resistant AML are preferentially reliant on the expression of MCL1 and consequently vulnerable to MCL1 genetic or pharmacologic perturbation^5,22,24,25^. Performing single cell RNA sequencing (sc-RNA-seq) on functionally validated ven/aza-sensitive and ven/aza-resistant primary AML specimens, we confirmed a 5-fold transcriptional upregulation of *MCL1* specific to ven/aza therapy resistance (Figure 1A, Supplemental Figure 1A). Furthermore, western blot analysis from representative primary ven/aza-sensitive and -resistant specimens highlights a significant 9-and 12-fold increase in MCL1 protein expression measured within the LSC fraction and bulk blast AML populations, respectively (Figure 1B-C and Supplemental Figure 1 B-C). Notably, and consistent with previous reports^5^, these ven/aza-resistant AML specimens also displayed a significant 5- and 10-fold reduction in BCL2 protein levels measured within LSCs and bulk AML, respectively (Figure 1B-C and Supplemental Figure 1 B-C). Thus, these data imply that a unique subset of primary ven/aza-resistant AML specimens display preferential expression of MCL1. Still, from a mechanistic standpoint, this preference for MCL1 among the BCL2 family homologs remains unexplained. To this end, we sought to further investigate the unique role of MCL1 for ven/aza resistance in AML.

**Figure 1.**
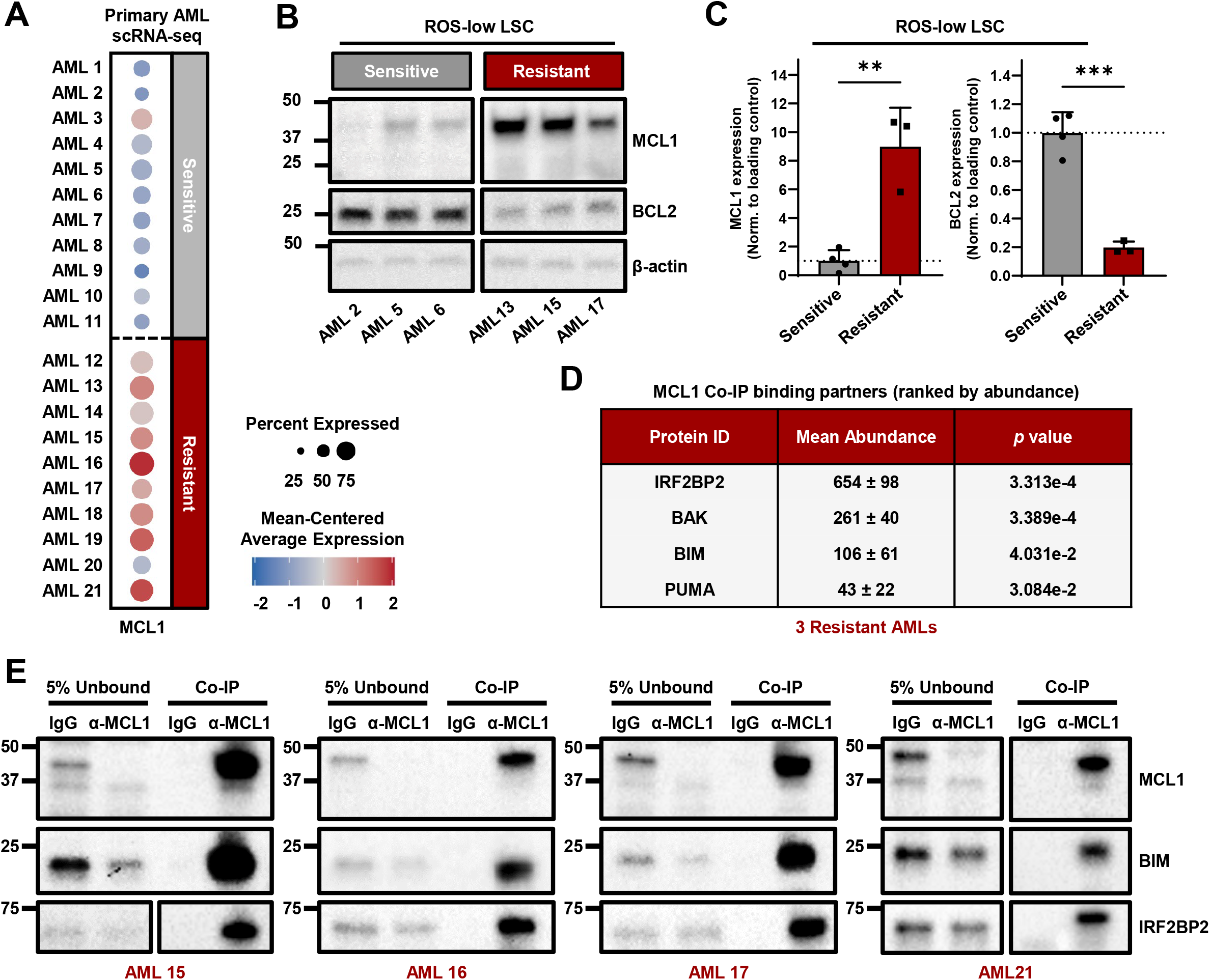
Ven/aza-resistant primary AML display preferential expression of MCL1 that interacts with the transcriptional repressor IRF2BP2. **(A)** Dot plot representing *MCL1* expression and proportion within an AML diseased cluster from scRNA-sequencing analysis of respective ven/aza-sensitive (N=11) and -resistant (N=10) primary AML samples. **(B)** Western blotting analysis for MCL1 and BCL2 protein expression within representative ven/aza-sensitive (N=3) and -resistant (N=3) primary AML ROS-low LSC. **(C)** Quantification of MCL1 and BCL2 protein expression within ven/aza-sensitive (N=4) and -resistant (N=3) primary AML ROS-low LSC. Data were normalized to β-actin loading control and presented as mean ± SD. Significance was determined using a two-tailed unpaired t test, ^**^p<0.01 and ^***^p<0.001. **(D)** Liquid chromatography-mass spectrometry analysis of MCL1 co-immunoprecipitated proteins within primary ven/aza-resistant AML (N=3). Depicted are mean ± SD abundance values from canonical apoptotic MCL1 binding partners as well as the most abundant precipitate, IRF2BP2. Significance was determined using a two-tailed unpaired t test and p values are indicated. **(E)** Western blotting validation of MCL1 co-immunoprecipitation within primary ven/aza-resistant AML (N=4). 5% of the unbound lysate was run for Isotype IgG and α-MCL1, compared against their respective bead capture and elution fractions (Co-IP). Membrane was probed for MCL1, BIM and IRF2BP2. See also Figure S1.

### MCL1 co-immunoprecipitates with the transcriptional repressor IRF2BP2 in ven/aza-resistant AML

To interrogate the underlying functional dependency for MCL1 within primary ven/aza-resistant AML, we performed co-immunoprecipitation (co-IP) of MCL1 coupled with mass spectrometry analysis (Figure 1D-E and Supplemental Figure 1D-F). We observed a near complete and robust MCL1 enrichment from ven/aza-resistant AML samples, evidenced by MCL1 expression that was exclusive to the anti-MCL1 co-IP fraction, as well as the absence of MCL1 protein within the unbound fraction subjected to the anti-MCL1 antibody (Figure 1E). Mass spectrometry analysis of the MCL1 immunoprecipitation revealed just over 100 significantly enriched proteins (Supplemental Figure 1E and Supplemental Table 1). Validating the success of MCL1 co-immunoprecipitation, canonical apoptotic and bona-fide MCL1 interacting partners like BAK, BIM and PUMA were observed in high abundance in ven/aza-resistant AML (Figure 1D and Supplemental Figure 1F). Gene Ontology and KEGG pathway analyses of the enriched proteins highlight involvement with an array of cellular processes such as programed cell death pathways, mitochondrial organization, ATP synthesis, oxidative phosphorylation, and very long-chain fatty acid metabolism (Supplemental Figure 1G-H). The apoptosis program shows enrichment with expected canonical proteins like BAK, BIM, BBC3/PUMA, and BMF, while the very long-chain fatty acid metabolism pathway displays enrichment of several rate limiting enzymes such as ACSL1, CPT1A and VLCAD which have been previously reported to functionally interact with MCL1 to control cellular metabolism^20,26^. However, the top MCL1-interacting protein by abundance (Figure 1D-E and Supplemental Table 1) revealed by mass spectrometry analysis was Interferon Regulatory Factor 2 Binding Protein 2 (IRF2BP2), a transcriptional repressor recently reported to be highly expressed and functionally relevant in AML blasts^27,28^. Importantly, the robust interaction observed between IRF2BP2 and MCL1 within ven/aza-resistant AML specimens is not observed (or shared) with BCL2^29^, suggesting a potentially unique function in supporting MCL1-driven therapy-resistance.

### IRF2BP2 transcriptional repressive activity segregates across ven/aza sensitivity in AML

Nuclear transcriptional repressive activity of IRF2BP2 has been reported to largely influence inflammatory and differentiation gene programs in various tissues^27,28,30-35^. In AML, Ellegast et al identified IRF2BP2-mediated repression of an inflammatory transcriptional signature (107 genes, largely NFκB via TNFα) through integrated chromatin immunoprecipitation sequencing (ChIP-seq) analysis and RNA-seq experiments. Proteolytic targeting of IRF2BP2 in AML cell lines and patient derived xenograft (PDX) models demonstrated strong activation of inflammatory genes, leading to potent cell death induction. Given the robust co-immunoprecipitation of IRF2BP2 with MCL1 within ven/aza-resistant primary AML, we sought to determine whether ven/aza sensitivity correlated with any IRF2BP2-dependent transcriptional changes. To understand the transcriptional impact that the MCL1-IRF2BP2 interaction has on ven/aza-resistant AML, we queried our primary AML scRNA-seq data set for IRF2BP2 transcriptional targets (107 genes^27^, denoted as IRF2BP2 signature) and found them to be overrepresented among ven/aza-resistant AML specimens (Figure 2A and Supplemental Figure 2A). Importantly, the IRF2BP2 signature displayed a strong correlation with *MCL1*, suggesting a potential loss of IRF2BP2 nuclear transcriptional repressive activity specific to ven/aza resistance (Figure 1A and Supplemental Figure 2B). Conversely, the inversion of the IRF2BP2 signature (denoted as IRF2BP2 signature-INV) shows a strong expression pattern with ven/aza-sensitive AML specimens, implying nuclear transcriptional repressive activity (Figure 2A and Supplemental Figure 2A). Notably, when we query an independent cohort of ven/aza-resistant patient samples, the collective transcriptional score of IRF2BP2 targets were overrepresented within nearly 75% of upfront refractory patients which also show higher levels of *MCL1* expression (Supplemental Figure 2C). Collectively, these data provide correlative evidence and clinical support for a potential link between *MCL1* expression, IRF2BP2 transcriptional regulation and overall response to venetoclax-based therapeutic regimens in AML.

**Figure 2.**
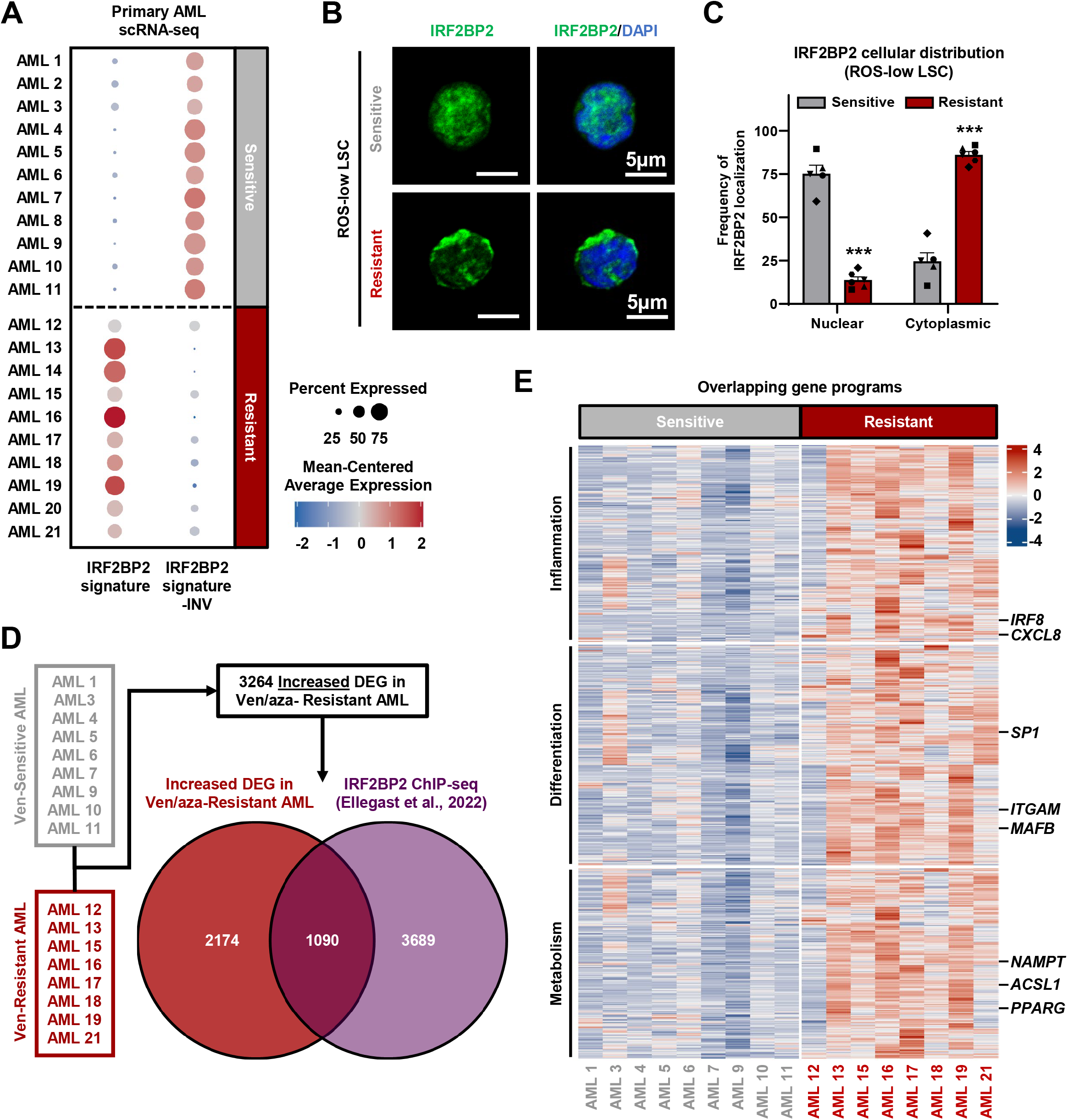
IRF2BP2 has differential nuclear/cytoplasmic localization and transcriptional repressive activity among ven/aza-sensitive and -resistant LSCs. **(A)** Dot plot representing expression and proportion for IRF2BP2 signature (107 genes, Ellegast et al., 2022) and IRF2BP2 signature-Inversion within an AML diseased cluster from scRNA-sequencing analysis of respective ven/aza-sensitive (N=11) and -resistant (N=10) primary AML samples. **(B)** Representative confocal microscopy analysis of IRF2BP2 expression and cellular localization within ven/aza-sensitive (N=5) and -resistant (N=6) primary AML ROS-low LSC. **(C)** Quantification of IRF2BP2 nuclear and cytoplasmic localization within ven/aza-sensitive (N=5) and -resistant (N=6) primary AML ROS-low LSC. Data were collected from an average of 100 individual cells per primary AML specimen and presented as a proportion with mean ± SD. Significance was determined using a two-tailed unpaired t test, ^***^p<0.001. **(D)** scRNA-seq differentially expressed gene (DEG) analysis between ven/aza-sensitive (N=9) and ven/aza-resistant (N=8) AML, resulting in 3264 significantly upregulated genes within ven/aza-resistant AML. Upregulated DEG in ven/aza-resistant AML were compared with an independent IRF2BP2 ChIP-seq dataset (Ellegast et al.,2022), revealing 1090 overlapping genes. **(E)** Heat map showing gene program annotations and expression of 665 overlapping genes (from 913 with gene program assignments). in ven/aza-sensitive (N = 9) and-resistant (N = 8) AML. *IRF8* and *CXCL8* are highlighted within the inflammation gene program (214 genes). *SP1, ITGAM* and *MAFB* are identified within a differentiation gene program (243 genes). *ACSL1, PPARG* and *NAMPT* are highlighted as essential metabolic regulators from the metabolism gene program (208 genes). See also Figure S2.

### IRF2BP2 is sequestered in the cytoplasm of ven/aza-resistant LSC, leading to the loss of nuclear transcriptional repression and subsequent overexpression of *ACSL1*

Nuclear localization of IRF2BP2 has been reported to be controlled by a specific post-translational modification (phosphorylation of serine 360) immediately downstream of its nuclear localization sequence^36^. Whole cell proteomics analysis of ven/aza-sensitive and -resistant LSC revealed reduced total IRF2BP2 as well as phosphorylation of IRF2BP2 at Serine 360 specific to ven/aza-resistant specimens (Supplemental Figure 2D-E), indicating a potential reduction in the nuclear accumulation of IRF2BP2, which could explain the overrepresented IRF2BP2 transcriptional score observed in our scRNA-seq analysis. Indeed, using confocal microscopy analysis, we observed that primary ven/aza-sensitive LSC display substantial nuclear IRF2BP2 localization while ven/aza-resistant LSC exhibit predominantly cytoplasmic IRF2BP2 expression (Figure 2B-C). To investigate the broader transcriptional consequences of MCL1-driven IRF2BP2 cytoplasmic sequestration, we overlapped the differentially upregulated genes within ven/aza-resistant AML from our scRNA-seq data set with putative IRF2BP2 transcriptional targets^27^ and identified 1090 significantly upregulated genes likely to result from the loss of nuclear IRF2BP2-mediated transcriptional repression (Figure 2D). Specifically, of the 1090 overlapping genes, nearly half (457 genes) align within known IRF2BP2 gene programs like inflammation or differentiation (Figure 2E). Additionally, another 20% (208 genes) of the overlapping gene-set categorizes within a metabolism-specific gene program (Figure 2E). Thus, the overlap between these two independent datasets, implies that IRF2BP2 may control critical components of several cellular processes unique to ven/aza-resistant AML pathology such as inflammation, differentiation and metabolism.

Notably, our group, as well as others, have implicated fatty acid metabolism (FAM) to be an essential process underlying therapy resistance in AML^6,21^. However, the underlying mechanisms that control activation of FAM in AML are unknown. Of particular interest, and functionally relevant to resistant AML metabolism, was an 11-fold increase in acyl-CoA synthetase long-chain family member 1 (*ACSL1*) transcript, an essential rate-limiting enzyme of FAM (Figure 2E and Supplemental Figure 2F). We validated the increased *ACSL1* transcript expression within ven/aza-resistant LSCs using quantitative PCR (Supplemental Figure 2G). In addition, other relevant metabolic regulators like Nicotinamide phosphoribosyltransferase (*NAMPT)*^10^, and Peroxisome proliferator-activated receptor gamma (*PPARG)*^*37*^, were also up-regulated in ven/aza-resistant specimens (Supplemental Figure 2G). Collectively, the data show that IRF2BP2 is aberrantly sequestered in the cytoplasm of ven/aza-resistant LSCs resulting in the loss of its nuclear transcriptional repressive activity and subsequent overexpression of metabolic targets like *ACSL1, NAMPT, and PPARG*.

### IRF2BP2 BH3-like α-helix motifs provide a functional domain-mediated interaction with MCL1 in ven/aza-resistant LSCs

Next, we wanted to determine if IRF2BP2 transcriptional regulation in AML is mediated through its cytoplasmic interaction with MCL1. The BCL2 family of proteins utilize their BCL2 homology 3 (BH3) domain as a key structural component to facilitate interactions amongst one another and to elicit their apoptotic and anti-apoptotic functions^38-40^. Additionally, many non-canonical functions of BH3-containing proteins are mediated through a cryptic or BH3-like motif within the interacting partner^20,26^. Indeed, BH3 motif analysis suggests that IRF2BP2 contains several BH3-like alpha helical domains^39,40^ (indicated with bold red sequence alignment) that likely mediate an interaction with the hydrophobic binding pocket of MCL1 (Figure 3A and Supplemental Figure 3A). To test the potential interaction between the BH3-like alpha helices of IRF2BP2 with MCL1, we performed an MCL1 co-IP from ven/aza-resistant AML following MCL1 BH3 disruption using the BH3-mimetic S63845^41^, a selective inhibitor that binds the hydrophobic BH3 motif binding pocket of MCL1. This resulted in a loss of IRF2BP2 co-immunoprecipitation, along with known BH3-interacting partner BIM, (Supplemental Figure 3B-C). Additionally, pharmacologic perturbation of the IRF2BP2-MCL1 interaction, using S63845, induces a rapid (1 hour) re-localization of IRF2BP2 from the cytoplasm into the nucleus of ven/aza-resistant LSC (Figure 3B-C). Importantly, this effect is not observed within ven/aza-sensitive LSCs that already display predominantly nuclear IRF2BP2 localization (Figure 3B-C).

**Figure 3.**
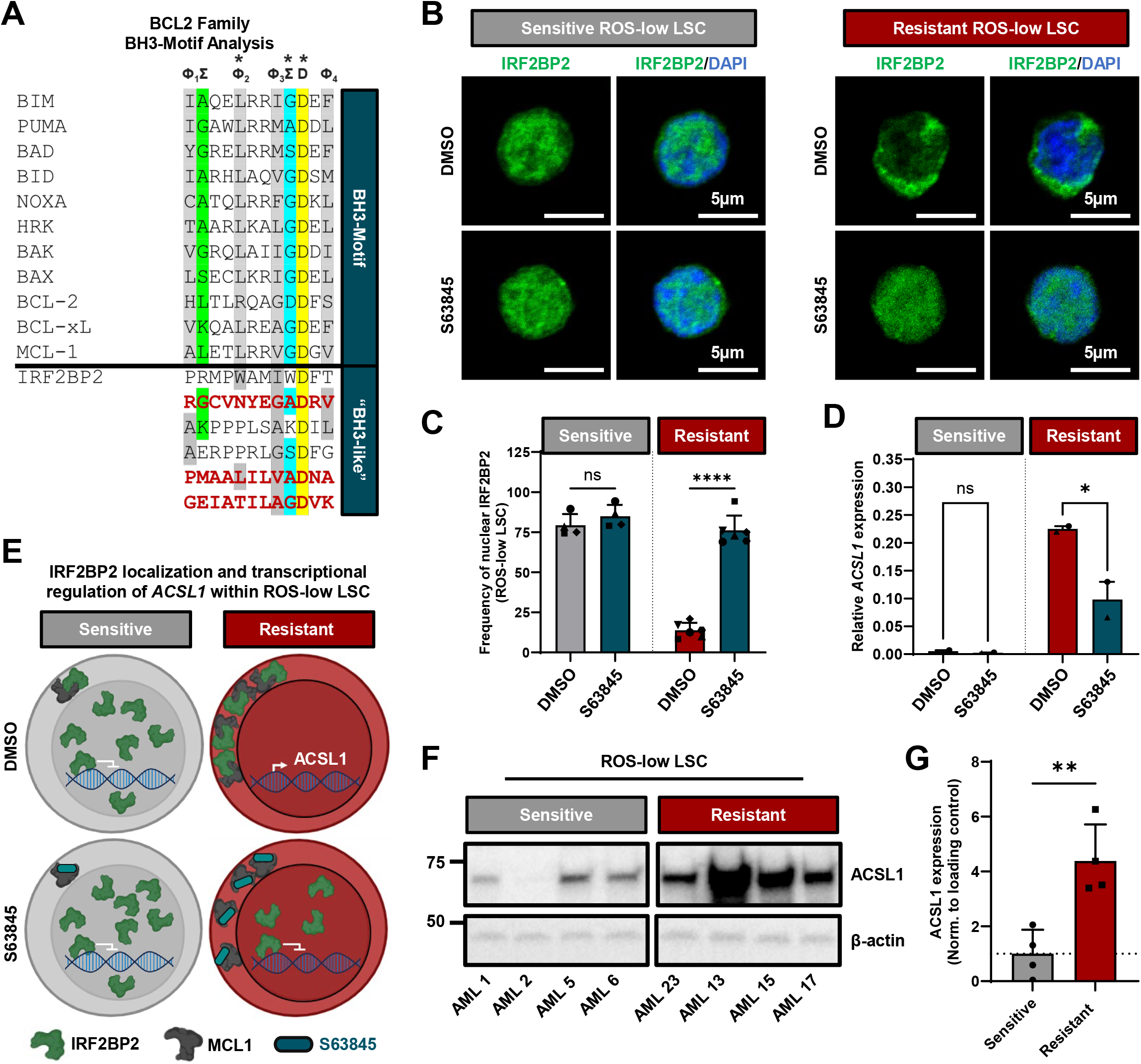
BH3-mimetic MCL1 inhibition redistributes IRF2BP2 into the nucleus of ven/aza-resistant LSCs and facilitates transcriptional repression on *ACSL1*. **(A)** BCL2 Family BH3-motif sequence alignment with IRF2BP2. The consensus sequence of BH3 domain (Aouacheria et al., 2015; Day et al., 2008) is depicted with Φ_1-4_ representing hydrophobic residues where Φ_2_ is usually a leucine (L), Σ representing small residues like glycine (G), alanine (A) and serine (S), and D representing the conserved aspartic acid residue. IRF2BP2 sequence homology predicts three BH3-like motifs that are in partial α-helical structure depicted in bold red font. **(B)** Confocal microscopy analysis of IRF2BP2 expression and cellular localization within representative ven/aza-sensitive (N=4) and -resistant (N=6) primary AML ROS-low LSC subjected to vehicle control or S63845 treatment for 1hr. **(C)** Quantification of IRF2BP2 nuclear localization within representative ven/aza-sensitive (N=4) and -resistant (N=6) primary AML ROS-low LSC subjected to vehicle control or S63845 treatment for 1hr. Data were collected from an average of 100 individual cells per primary AML specimen from each treatment condition and presented as a proportion with mean ± SD. Significance was determined using a two-way ANOVA test, ^****^p<0.0001. **(D)** Relative *ACSL1* mRNA expression in representative ven/aza-sensitive (N=2) and -resistant (N=2) primary AML ROS-low LSC subjected to vehicle control or S63845 treatment for 4hr. Data are presented as mean ± SD relative to gene loading control. Significance was determined using a two-way ANOVA test, ^*^p<0.05. **(E)** Schematic representing IRF2BP2 cellular localization and transcriptional repressive activity on *ACSL1* within ven/aza-sensitive and -resistant primary AML ROS-low LSC subjected to vehicle control or S63845. **(F)** Western blotting analysis for ACSL1 protein expression within representative ven/aza-sensitive (N=4) and -resistant (N=4) primary AML ROS-low LSC. **(G)** Quantification of ACSL1 protein expression within ven/aza-sensitive (N=4) and -resistant (N=4) primary AML ROS-low LSC. Data were normalized to β-actin loading control and presented as mean ± SD. Significance was determined using a two-tailed unpaired t test, ^**^p<0.01. See also Figure S3.

Similarly, *ACSL1* transcript expression is exclusively observed within ven/aza-resistant LSCs where cytoplasmic IRF2BP2 localization and a functional loss of nuclear transcriptional repressive activity were detected (Figure 3D). Conversely, IRF2BP2 nuclear re-localization, using BH3-mimetic S63845, resulted in a rapid transcriptional repression of *ACSL1*, as well as other IRF2BP2 putative targets like *IL1B* and *NAMPT* (Figure 3D and Supplemental Figure 3D). Notably, transcripts were measured after only 4 hours of exposure to S63845 as MCL1 inhibition is extremely cytotoxic to these cells. These data clearly indicate a role for IRF2BP2 in repressing genes like *ACSL1, NAMPT* and *IL1B* when it is localized in the nucleus. Notably, we observe some AML patient specimens bearing both ven/aza-sensitive and -resistant subpopulations. Analysis of intra tumoral heterogeneity in these patients demonstrates IRF2BP2 nuclear re-localization and transcriptional repression of IRF2BP2 targets (*ACSL1, NAMPT* and *IL1B*) upon treatment with an MCL1 inhibitor that is exclusive to the ven/aza-resistant LSC population (Supplemental Figure 3E).

Collectively, MCL1-directed IRF2BP2 localization within ven/aza-sensitive and -resistant LSCs dictates subsequent transcriptional output of metabolic genes like *ACSL1*. Nuclear IRF2BP2 in ven/aza-sensitive LSCs represses *ACSL1*, while cytoplasmic MCL1-IRF2BP2 in ven/aza-resistant LSCs results in upregulated *ACSL1* expression. Furthermore, MCL1-IRF2BP2 disruption within ven/aza-resistant LSCs, using BH3-mimetic S63845, results in IRF2BP2 nuclear accumulation and transcriptional repression activity (Figure 3E). In line with increased *ACSL1* transcript, whole cell proteomics and western blot analyses confirmed elevated ACSL1 protein levels, suggesting a potential role for ACSL1 in controlling FAO within ven/aza-resistant LSCs (Figure 3F-G and Supplemental Figure 3F).These data highlight a unique MCL1-driven loss of IRF2BP2 transcriptional repressive activity that defines ven/aza-resistant AML pathology, wherein genes involved in fatty acid metabolism are activated.

### ACSL1 promotes FAM in ven/aza-resistant LSCs

FAO regulation by MCL1 has been shown as independent of its anti-apoptotic activity, and MCL1 has been labeled as a master regulator controlling multiple levels of FAO in human cancers (transcriptional, protein and subsequent metabolites)^42^. The underlying mechanisms regulating such FAO machinery and control is not well understood. Our data suggests that MCL1 is functioning within LSCs to sequester IRF2BP2 in the cytoplasm, releasing the transcriptional repression of crucial FAO enzymes like *ACSL1*.

The ACSL enzymatic family converts free long-chain fatty acids into fatty acyl-CoA esters, and thereby plays a key role in fatty acid degradation as well as lipid synthesis^43,44^. Of the five ACSL family members (ACSL1 and ACSL3-6), *ACSL1, ACSL3* and *ACSL4* were observed to be upregulated in ven/aza-resistant AML through our scRNA-seq analysis (transcript fold change increase over ven/aza-sensitive AML of 11.3, 1.6 and 1.9, respectively). However, *ACSL1* had the highest fold change over ven/aza-sensitive AML and was the only family member that overlapped with putative IRF2BP2 transcriptional targets (Figure 2E). ACSL1 has a broad substrate specificity for long-chain fatty acids (FA), specifically 16 to 18 carbon saturated FAs and 16 to 20 carbon unsaturated FAs^45^. Global metabolomic analysis between ven/aza-sensitive and ven/aza-resistant LSCs identified a significant accumulation of long-chain FAs, both saturated and unsaturated, as well as long-chain acyl-carnitines and 18 carbon sphingosine within ven/aza-resistant LSCs, at the expense of depleted amino acid metabolites (Figure 4A and Supplemental Figure 4A-C). Specifically, long-chain acyl-carnitine species like palmitoyl carnitine (16:0), 9-hexadecenoyl carnitine (16:1), oleoyl carnitine (18:1), linoleyl carnitine (18:2), and arachidonoyl carnitine (20:4), as well as FAs like linoleic acid (18:2) and arachidonic acid (20:4) were significantly increased within ven/aza-resistant LSCs (Figure 4A and Supplemental Figure 4B-C). Importantly, CPT1A can subsequently convert ACSL1-derived acyl-CoA species into acyl-carnitines for subsequent β-oxidation^46^. Moreover, linoleic acid metabolism (FA 18:2) was the top enriched pathway represented by the metabolite changes between ven/aza-sensitive and -resistant LSCs (Figure 4B). Notably, the significantly upregulated acyl-carnitine species observed in ven/aza-resistant LSCs are likely direct products derived from long-chain FAs with substrate specificity for ACSL1.

**Figure 4.**
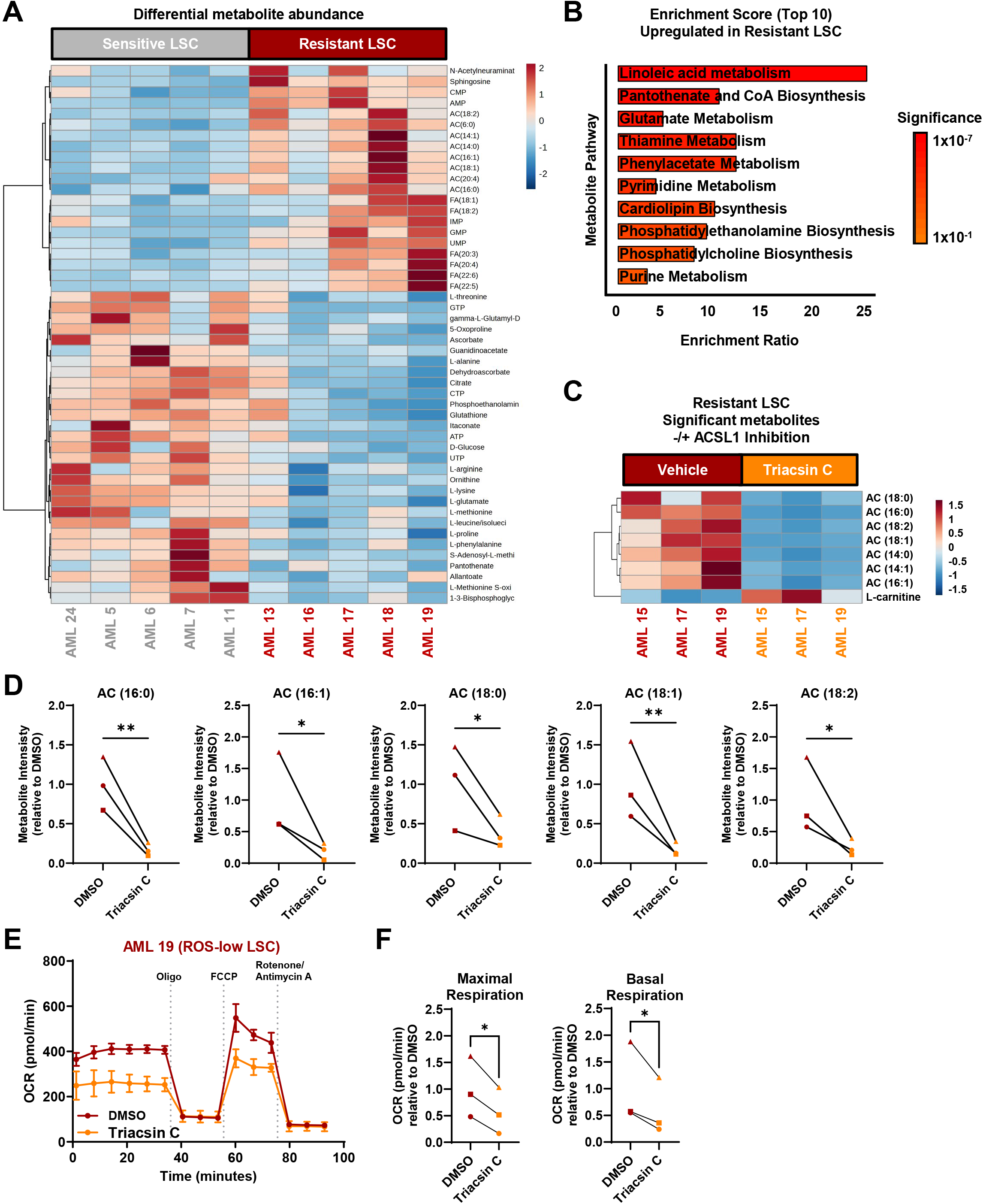
Ven/aza-resistant LSCs rely on an abundance of long-chain FA and ACSL1 to fuel β-oxidation. **(A)**Heat map depicting the top 50 significant metabolites by p-value between ven/aza-sensitive (N=5) and - resistant (N=5) primary AML ROS-low LSC. **(B)** Metabolite pathway enrichment analysis representing the top 10 upregulated metabolic pathways in ven/aza-resistant primary AML ROS-low LSC. **(C)** Heat map depicting significant metabolite changes within ven/aza-resistant (N=3) primary AML ROS-low LSC subjected to vehicle control or Triacsin C treatment for 16 hours. **(D)** Long-chain acyl-carnitine abundance within ven/aza-resistant (N=3) primary AML ROS-low LSC subjected to vehicle control or Triacsin C treatment for 16hr. Significance was determined using a two-tailed ratio paired t test, ^*^p<0.05, ^**^p<0.01. **(E)** Representative oxygen consumption rate (OCR) curve from Seahorse XF Palmitate Oxidation Stress assay comparing respiration of ven/aza-resistant primary AML ROS-low LSC subjected to vehicle control or Triacsin C treatment for 16 hours. Technical replicates of four per data point. Data are presented as mean ± SD for 100k cells. Vertical dotted lines indicate injection times used throughout assay. **(F)** Maximal and basal respiration rates calculated from Seahorse XF Palmitate Oxidation Stress assay from ven/aza-resistant (N=3) primary AML ROS-low LSC subjected to vehicle control or Triacsin C treatment for 16 hours. Data are presented as OCR relative to DMSO. Significance was determined using a two-tailed ratio paired t test, ^*^p<0.05. See also Figure S4.

To test whether ACSL1 was primarily responsible for the increased abundance of long-chain acyl-carnitine species observed within ven/aza-resistant LSCs, we performed metabolomic analysis on ven/aza-resistant LSCs following ACSL1 pharmacologic inhibition using Triacsin C^26,47^. Inhibition of ACSL1 within ven/aza-resistant LSCs resulted in a very specific depletion of long-chain acyl-carnitines highlighting a potential role for ACSL1 in coordinating FAO within therapy resistant AML (Figure 4C-D and Supplemental Figure 4D). We have previously reported that monocytic AML, which strongly correlate with ven/aza-resistance, display elevated basal respiration rates and oxidative phosphorylation compared to more primitive and ven/aza-sensitive AML^5^. Our current data suggest that the observed increase in respiration is likely the result of switching metabolic input from primarily amino acid metabolism^8^ (Figure 4A and Supplemental Figure 4B) toward an ACSL1-dependent FA β-oxidation route. Seahorse analysis, specifically measuring long-chain FAO of palmitate (16:0) via oxygen consumption, confirmed ACSL1 is metabolically essential for ven/aza-resistant LSC basal and maximal respiration rates (Figure 4E-F). Importantly, Triacsin C reduced maximal respiration rates with comparable levels to CPT1A inhibition/ etomoxir treatment (Supplemental Figure 4E-F) when palmitate is added to the assay media as a substrate (Seahorse XF Palmitate Oxidation Stress Test Kit). These data confirm ACSL1 is exclusively upregulated within ven/aza-resistant AML LSCs and uniquely drives the conversion of long-chain FAs to intermediate fatty acyl-CoAs, which are subsequently transferred into the mitochondria as acyl-carnitines for FAO.

### Pharmacologic inhibition of ACSL1 functionally impairs ven/aza-resistant AML

Our previous studies have demonstrated that activation of FAO acts to circumvent the mechanism by which ven/aza treatment kills LSCs^6,8^. Thus, we sought to determine whether targeting ACSL1 could functionally impair primary ven/aza-resistant LSCs. Our data show that pharmacologic inhibition of ACSL1 (using Triacsin C) (Figure 5A), selectively reduces the viability of ven/aza-resistant AML specimens (Figure 5B and Supplemental Figure 5A). Notably, while the ven/aza-sensitive specimens show a significant response to ven/aza therapy, no significant response to ACSL1 inhibition was detected (Figure 5B and Supplemental Figure 5A), likely due to the reduced abundance of FAs and ACSL1 protein expression (Figure 4A and Figure 3F-G). Additionally, ven/aza-resistant AML specimens display an exacerbated 80% reduction in colony forming potential following ACSL1 inhibition (Figure 5C and supplemental figure 5B). Importantly, we observed no change in colony forming potential of age matched normal CD34+ stem and progenitor cells isolated from BM and subjected to overnight ACSL1 inhibition/ Triacsin C treatment (Figure 5C and Supplemental Figure 5B). Consistent with the observed *in vitro* functional deficiencies, ven/aza-resistant AML exhibited a significant loss of engraftment potential in immune-deficient mice following treatment with Triacsin C (Figure 5D). Together, these data support the hypothesis that ven/aza-resistant AML are functionally reliant on FAO as they are uniquely vulnerable to pharmacologic perturbation of ACSL1.

**Figure 5.**
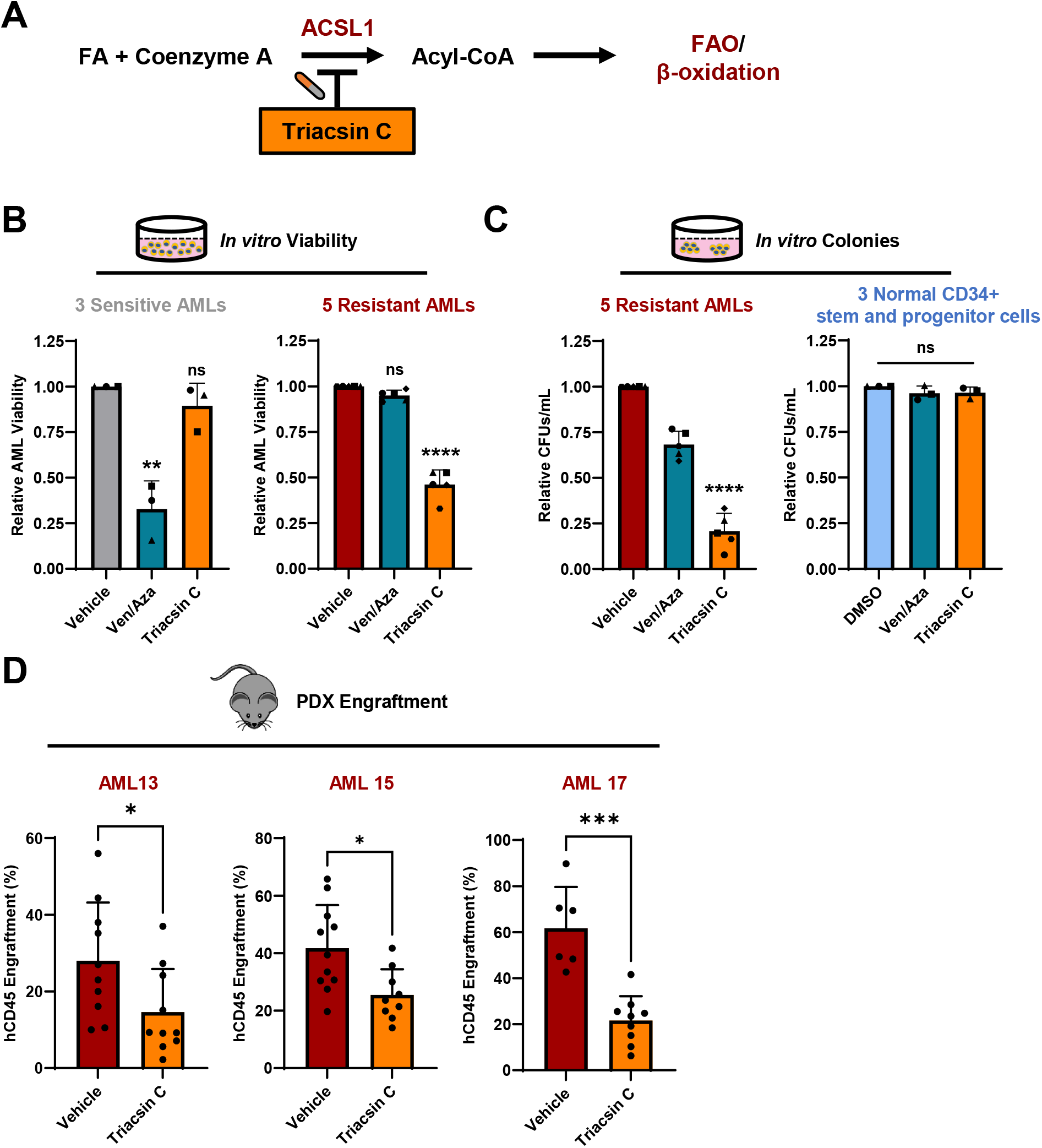
Ven/aza-resistant AML are functionally reliant on ACSL1. **(A)** Schematic representing ACSL1 enzymatic activity. ACSL1 drives the conversion of free long-chain fatty acids into fatty acyl-CoA species, a common precursor to acyl-carnitine-mediated mitochondrial entry and fatty acid β-oxidation. **(B)** *In vitro* viability of ven/aza-sensitive (N=3) and -resistant (N=5) primary AML subjected to ven/aza or Triacsin C treatment for 24 hours. Data are presented as mean ± SD. Significance was determined using two-tailed unpaired t test, ^**^p<0.01, ^***^p<0.001, ^****^p<0.0001. **(C)** *In vitro* colony forming potential of ven/aza-resistant (N=5) primary AML and normal CD34+ hematopoietic stem and progenitor cells (N=3) subjected to ven/aza or Triacsin C treatment for 16 hours. Data are presented as mean ± SD. Significance was determined using two-tailed unpaired t test, ^****^p<0.0001. **(D)** *Ex vivo* engraftment potential of ven/aza-resistant (N=3) primary AML following Triacsin C treatment for 16 hours. Data are presented as mean ± SD. Significance was determined using a two-tailed unpaired t test, ^*^p<0.05, ^***^p<0.001. See also Figure S5.

## Discussion

The existence and persistence of LSCs following venetoclax-based therapies continues to be a significant clinical challenge. Both upfront resistance as well as relapse following an initial response demonstrates an unmet need to better understand the underlying resistance mechanisms within LSC subtypes. Our data provide evidence that MCL1 expression in ven/aza-resistant LSCs drives IRF2BP2 cytoplasmic sequestration and consequent activation of *ACSL1*, promoting fatty acid oxidation in therapy-resistant primary AML (Figure 6A). Pharmacological BH3 motif interference, using MCL1-specific S63845, resulted in MCL1-IRF2BP2 dissociation followed by rapid nuclear redistribution of IRF2BP2 and transcriptional repression of target genes like *ACSL1*. Notably, BH3-mimetics like S63845 will eradicate ven/aza-resistant AML likely due to both canonical apoptotic and non-canonical metabolic-dependent functions of MCL1. However, clinical use of several MCL1 BH3-mimetic inhibitors has demonstrated dose-limiting cardiac toxicities indicating this approach may not be feasible for patients^48-50^. Alternatively, our findings show that pharmacological inhibition of ACSL1, downstream of the MCL1-IRF2BP2 axis, severely limits the ability of ven/aza-resistant LSCs to catabolize FAs, resulting in a selective and targetable vulnerability in therapy-resistant AML specimens (Figure 6B). Further, inhibition of ACSL1 was sufficient to significantly impair LSC engraftment activity. Thus, we posit that ACSL1 represents a key component of the overall mechanism underlying ven/aza resistance within LSCs. Ongoing studies will further dissect the IRF2BP2 target genes/pathways that mediate venetoclax resistance. Indeed, the IRF2BP2 transcriptional repressive activity mediates several pathways such as inflammation, differentiation, and metabolism, which may augment the activity of ACSL1 in driving venetoclax resistance in AML.

**Figure 6.**
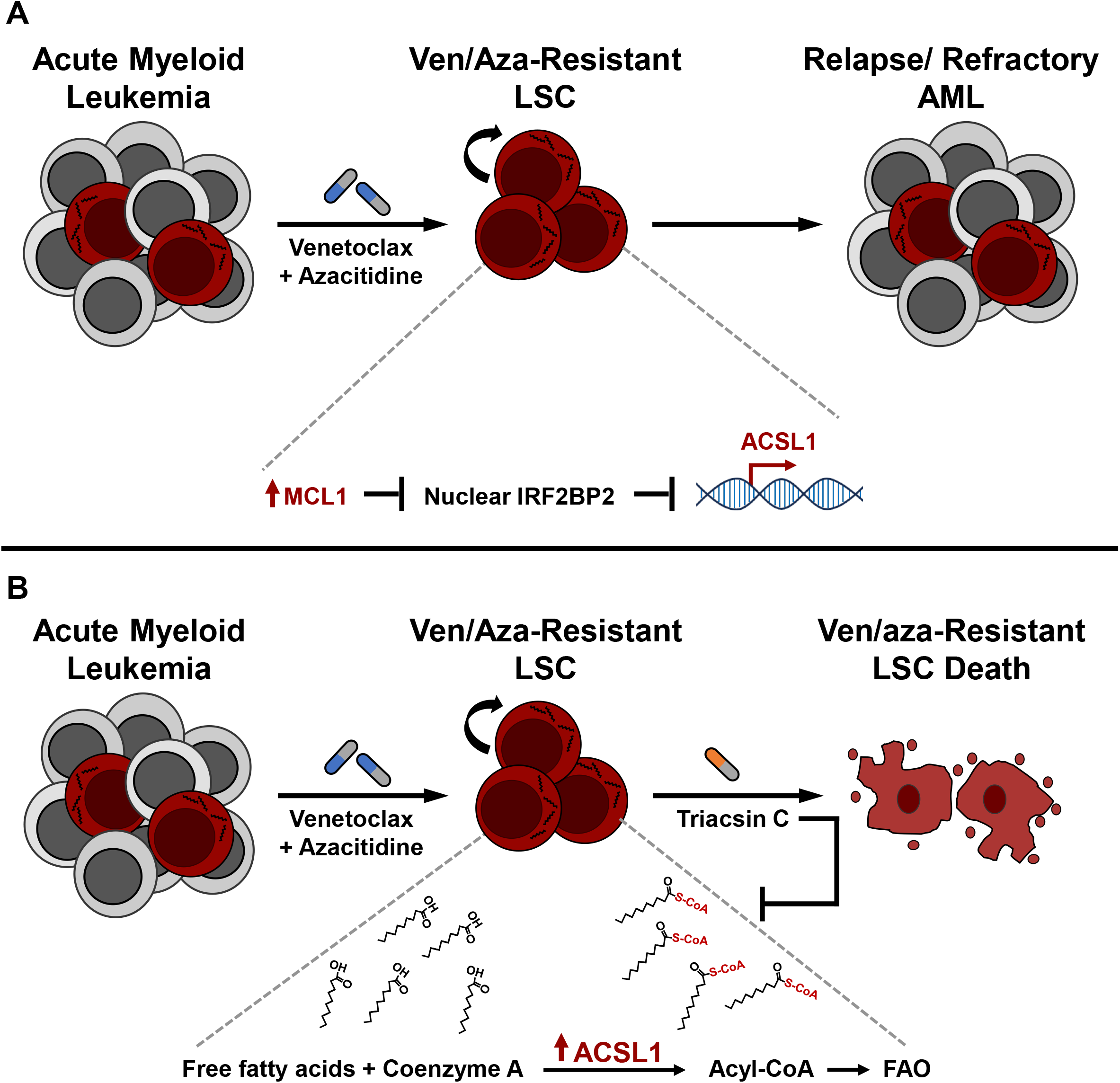
Working Model/ schematic. **(A)** Novel mechanism by which MCL1 non-canonically drives IRF2BP2 cytoplasmic sequestration and inhibition of transcriptional repressive activity uniquely within ven/aza-resistant primary AML ROS-low LSC, resulting in the subsequent upregulation of a metabolic program including *ACSL1*. ACSL1 drives the conversion of free fatty acids into fatty acyl-CoA species, a common precursor to acyl-carnitine-mediated mitochondrial entry and fatty acid β-oxidation. Consequently, ven/aza-resistant primary AML are vulnerable to ACSL1/ fatty acid oxidation inhibition.

A role for IRF2BP2 in transcriptionally repressing inflammation has been previously reported and recently highlighted in AML^27,28^. IRF2BP2 genetic or proteolytic depletion in AML was shown to unleash a cell-intrinsic inflammatory-induced mechanism of cell death^27^. Notably, these studies employed models in which IRF2BP2 is predominantly nuclear localized, and thereby acting as a repressor of genes associated with inflammation. Thus, the loss of IRF2BP2 resulted in a dramatic upregulation of IL1β/TNFα/NFκB transcripts and signaling^27^. Adding to these data, we provide substantial evidence demonstrating that AML LSCs exist in one of two distinct phenotypic states, with either nuclear or cytoplasmic localization of IRF2BP2. LSCs with nuclear IRF2BP2 would be inherently susceptible to the acute loss of its transcriptional repressive activities and thus represent an interesting context-specific therapeutic target^27^, whereas LSCs that naturally sequester IRF2BP2 in the cytoplasm appear to require IRF2BP2 target genes. Moreover, we showed a negative correlation between the IRF2BP2 signature (indicating loss of IRF2BP2 transcriptional repressive activity) and AML samples refractory to ven/aza therapy. Thus, interrogating IRF2BP2 localization, or a refined transcriptional signature within primary samples, may offer predictive benefits to patients with AML regarding treatment strategies.

We report significant transcriptional changes between ven/aza-sensitive and -resistant AML that likely, at least in part, results from the unique MCL1-mediated sequestration of IRF2BP2 within the cytoplasm of ven/aza-resistant LSCs. Notably, ven/aza-resistant AML often presents with monocytic differentiation features^5,51^ in addition to increased inflammatory signaling^5^ that may parallel a loss of nuclear IRF2BP2 activity. Indeed, suppressed differentiation is another consequence of IRF2BP2 transcriptional repression^28,31-35^. Along these lines, ven/aza-resistant AML, where IRF2BP2 is sequestered in the cytoplasm, show significant upregulation of traditional monocytic differentiated markers that overlap with IRF2BP2 ChIP-seq datasets such as *ITGAM, CD14*, and *CD68*. In addition, major myeloid-driving transcription factors/programs, like *MAFB*^52^, *SP1*^53,54^, *JUN/FOS*^55^ and *AHR/ARNT*^56^ appear to be under IRF2BP2 transcriptional control, and de-repressed in ven/aza-resistant LSCs. In normal hematopoiesis, *SPI1*/PU.1 drives monocytic differentiation by positively regulating the AP-1 transcription factors, JUN/FOS, and their expression highly correlates with M4/M5 monocytic AML subtypes^55^. Interestingly, *SP1*/*JUN*/*FOS* are all putative repressed IRF2BP2 targets^27^ and consequently upregulated in our ven/aza-resistant AML which displays cytoplasmic IRF2BP2 localization and loss of nuclear transcriptional repressive activity. Therefore, we propose that MCL1-mediated IRF2BP2 cytoplasmic sequestration may be a central regulator driving the monocytic phenotype in AML.

The understanding of lipid metabolism in cancer biology has expanded rapidly in recent years and has been implicated to support LSC function and AML disease progression through diverse mechanisms that extend beyond basic anabolism and catabolism of lipid species. Specifically, lipids have been implicated in supporting energy production, contributing toward membrane integrity and composition, as well as coordinating various signaling pathways and ferroptosis ^21,46,57-64^. Moreover, recent work has associated metabolic plasticity and lipid rewiring to underscore therapy response in AML^61,63^. Our work suggests that ACSL1, downstream of MCL1-IRF2BP2, is promoting long-chain FAO and venetoclax resistance in AML. Interestingly, the ACSL enzyme family convert free long-chain fatty acids into fatty acyl-CoA esters, and thereby play a key role in fatty acid degradation as well as lipid biosynthesis. Notably, acyl-CoA-binding proteins successfully transport acyl-CoAs to different metabolic sites within cells which can dictate their metabolism. Specifically, peroxisomes will utilize acyl-CoAs for both oxidation^65^ and synthesis of alkyl lipids^66^, whereas mitochondria primarily facilitate FAO, and the endoplasmic reticulum favors lipid synthesis^67^. Our work describes a role for ACSL1 in promoting FAO in ven/aza-resistant LSCs, However, additional studies are ongoing to explore a role for ACSL1 in mediating lipid synthesis, storage, and signaling within primary ven/aza-resistant AML.

In summary, these data demonstrate that MCL1 functions to control IRF2BP2 localization and thus alters its effects on transcriptional targets, resulting in LSC metabolism that hinders treatment response within AML. We propose that the transcriptional changes produced by the MCL1-IRF2BP2 cytoplasmic interaction provide a unique framework for dissecting venetoclax resistance mechanisms in AML.

## Resource availability

### Lead contact

Further information and requests for materials should be directed to the lead contact, Craig T. Jordan.

### Materials availability

This study did not generate new unique reagents.

### Data availability

The data that support the findings of this study are available from the corresponding author upon reasonable request. ScRNA-seq data is available in GEO (GSE232559).

## Acknowledgments

We thank all the patients and their families for consenting for tissue banking and use of primary AML samples for these studies. M.J.A. was supported by an NCI NRSA F32 training award (F32CA275350). C.T.J. is generously supported by the Nancy Carroll Allen Chair in Hematology Research, Leukemia and Lymphoma Society SCOR grants (7020-19 and 7033-24), NIH R01CA292071, and NIH R35CA242376. A.V. was supported by the Leukemia and Lymphoma Society Fellow award (5677-25). JME is supported by an Independent Investigator Award from the Rally Foundation for Childhood Cancer Research, a Fellow to Faculty Award from the American Society of Hematology, and a Physician-Scientist Research Grant from the European Hematology Association. We would also like to acknowledge the Colorado Cancer Center and Colorado Nutrition Obesity Research Center (NIH P30 DK48520) for providing the core for seahorse assays. Schematics were created in part through Biorender (https://BioRender.com).

## Author contributions

M.J.A. and C.T.J. designed studies; M.J.A., M.M., S.B.P., I.S., R.M., A.V., A.K., T.Y., M.D., D.S., and A.T. performed experiments; M.J.A., M.M., A.E.G., S.G., S.B.P., M.D., D.S., A.T., and A.D. performed data analysis; B.M.S., A.E.G., W.S., J.M.E, T.W., K.S., J.T.O., A.D., and C.A.S. provided critical reagents, insightful views and critiques; M.J.A. and C.T.J wrote the manuscript; all authors revised and approved manuscript.

## Supplemental information titles and legends

### Supplemental information

Document S1. Figures S1-S5

Document S2. Figure legends for Figures S1-S5.

Table S1. Top MCL1 Co-immunoprecipitated features ranked by abundance score.

Table S2. File containing primary AML information including age, sex, cytogenetics, and mutation information.

## Materials and Methods

### Human Specimens

Primary human AML specimens were obtained by apheresis product, peripheral blood, or bone marrow. Mobilized peripheral blood was obtained from normal healthy volunteer donors at the University of Colorado. All patients gave written informed consent for procurement of samples on the University of Colorado tissue procurement protocol (Colorado Multiple Institutional Review Board Protocol; #12-0173 and #06-0720). The University of Colorado Institutional Review Board approved the retrospective analysis. All specimens were acquired in accordance with recognized ethical guidelines (Declaration of Helsinki and U.S. Common Rule). Further details on each human AML specimen used for analysis are included in Supplemental Table S2 including age, sex, cytogenetics, and mutation information. Primary human AML specimens have been systematically evaluated for multiple biological properties, including ven/aza responsiveness. AML specimens indicated as sensitive or resistant to ven/aza in the following figures have been experimentally verified as such, with *in vitro* and *in vivo* data.

### Cell Sorting

Primary human AML specimens were sorted for ROS-low leukemia stem cells (LSCs) as previously described^7,68^. Briefly, specimens were thawed and stained with 4′,6-diamidino-2-phenylindole (DAPI; EMD Millipore, 278298; dilution 500 nmol/L) to exclude dead cells, CD19 (BD, 555413; dilution 1:20) and CD3 (BD, 557749; dilution 1:40) to exclude lymphocytes, CD45 (BD, 571875; dilution 1:40) to identify the blast population and CellROX deep red (Thermo Fisher, C10422; dilution 5 μmol/L) to identify the 20% of AML blasts with the lowest ROS stain signal, deemed “ROS-low LSCs”.

### MCL1 Co-Immunoprecipitation and Liquid chromatography-Mass Spectrometry (LC-MS)

#### Co-Immunoprecipitation

Co-Immunoprecipitations were performed using the Pierce Classic Magnetic IP/Co-IP Kit (Thermo Scientific, cat. #88804). Briefly, primary AML cells were lysed in Pierce IP lysis/wash reagent, supplemented with proteinase inhibitor (Thermo Scientific, 1862209), according to manufacturer’s instructions. Subsequently, 1mg of protein lysate was incubated with 1.5μg of either anti-MCL1 (Cell Signaling Technologies, 94296) or isotype control (Cell signaling Technologies, 3900) antibody overnight at 4°C. Antigen/antibody complex was then incubated with Pierce Protein A/G Magnetic Beads for 1.5 hours at room temperature. The beads were bound to a magnet and washed using Pierce IP lysis/wash reagent.

Antigen/antibody complex was eluted from the magnetic beads using Non-reducing Lane Marker Sample Buffer (Thermo Scientific, 1859594) with 50mM Dithiothreitol (DTT: Sigma, 43816).

#### Liquid chromatography-Mass Spectrometry (LC-MS)

Eluted samples were loaded onto a 1.5 mm thick NuPAGE Bis-Tris 4−12% gradient gel (Invitrogen). The gel was stained using SimplyBlue™ SafeStain (Invitrogen, Carlsbad, CA) and de-stained with water according to the manufacturer’s protocol. Each lane of the gel was divided into 6 equal-sized bands, and proteins in the gel were digested as follows. The pieces were destained in 200 µL of 25mM ammonium bicarbonate in 50% v/v acetonitrile for 15 minutes and washed with 200µL of 50% (v/v) acetonitrile. Disulfide bonds in proteins were reduced by incubation in 10mM dithiothreitol (DTT) at 60 °C for 30 minutes and cysteine residues were alkylated with 20mM iodoacetamide (IAA) in the dark at room temperature for 30 minutes. Gel pieces were subsequently washed with 100µL of distilled water followed by addition of 100mL of acetonitrile and dried on SpeedVac (Savant ThermoFisher). Then, 100ng of trypsin was added to each sample and allowed to rehydrate the gel plugs at 4°C for 45 minutes and then incubated at 37°C overnight. The tryptic mixtures were acidified with formic acid up to a final concentration of 1%. Peptides were extracted two times from the gel plugs using 1% formic acid in 50% acetonitrile. The collected extractions were pooled with the initial digestion supernatant and dried on SpeedVac (Savant ThermoFisher). Samples were desalted on Thermo Scientific Pierce C18 Tip.

#### Mass spectrometry analysis

20μL of each sample was loaded onto individual Evotips for desalting and then washed with 20μL 0.1% FA followed by the addition of 100μL storage solvent (0.1% FA) to keep the Evotips wet until analysis. The Evosep One system (Evosep, Odense, Denmark) was used to separate peptides on a Pepsep column, (150 μm inter diameter, 15cm) packed with ReproSil C18 1.9 μm, 120A resin. The system was coupled to the timsTOF Pro mass spectrometer (Bruker Daltonics, Bremen, Germany) via the nano-electrospray ion source (Captive Spray, Bruker Daltonics). The mass spectrometer was operated in PASEF mode. The ramp time was set to 100ms and 10 PASEF MS/MS scans per topN acquisition cycle were acquired. MS and MS/MS spectra were recorded from m/z 100 to 1700. The ion mobility was scanned from 0.7 to 1.50□Vs/cm^2^. Precursors for data-dependent acquisition were isolated within□±1□Th and fragmented with an ion mobility-dependent collision energy, which was linearly increased from 20 to 59□eV in positive mode. Low-abundance precursor ions with an intensity above a threshold of 500 counts but below a target value of 20000 counts were repeatedly scheduled and otherwise dynamically excluded for 0.4□minutes.

#### Database Searching and Protein Identification

MS/MS spectra were extracted from raw data files and converted into .mgf files using MS Convert (ProteoWizard, Ver. 3.0). Peptide spectral matching was performed with Mascot (Ver. 2.5) against the Uniprot mouse database. Mass tolerances were +/-15 ppm for parent ions, and +/-0.4 Da for fragment ions. Trypsin specificity was used, allowing for 1 missed cleavage. Met oxidation, protein N-terminal acetylation, peptide N-terminal pyroglutamic acid formation were set as variable modifications with Cys carbamidomethylation set as a fixed modification. Scaffold (version 5.01, Proteome Software, Portland, OR, USA) was used to validate MS/MS based peptide and protein identifications. Peptide identifications were accepted if they could be established at greater than 95.0% probability as specified by the Peptide Prophet algorithm. Protein identifications were accepted if they could be established at greater than 99.0% probability and contained at least two identified unique peptides.

### Immunoblotting

Cells were lysed using Pierce IP lysis/wash reagent (Thermo Scientific, cat. #88804) supplemented with proteinase inhibitor (Thermo Scientific, 1862209) according to manufacturer’s instructions. Lysates were subsequently denatured using 50mM DTT (DTT: Sigma, 43816) followed by boiling for 10 minutes. Protein lysates were loaded into a polyacrylamide gel and transferred to a polyvinylidene difluoride (PVDF) membrane using the mini trans-blot transfer system (Bio-Rad). Membranes were probed with primary antibodies of the following targets: MCL1 (Cell Signaling Technologies, no. 5453), BCL-2 (Cell Signaling Technologies, no.15071), BIM (Cell Signaling Technologies, 2933), β-actin (Cell Signaling Technologies, no.4970), Vinculin (Cell Signaling Technologies, 13901), ACSL1 (Cell Signaling Technologies, 9189), and IRF2BP2 (Proteintech, 18847-1-AP). Membranes were incubated with primary antibodies overnight at 4°C on a shaker. All primary antibodies were used at a 1:1,000 dilution. After overnight incubation, membranes were incubated with respective horseradish peroxidase-conjugated secondary antibodies (Bio-Rad) for 1 hour at room temperature. Pierce Clean Blot Detection reagent (Thermo Scientific, 21230) was used to probe immunoprecipitation blots for 1 hour at room temperature where denatured heavy- and light-chain IgG bands occlude protein of interest (close molecular weight to 50 or 25 kDa). Subsequently, blots were exposed to SuperSignal West Femto Maximum Sensitivity Substrate (Thermo Scientific, 34096) and imaged using the ChemiDoc Imaging System (Bio-Rad). Images were processed using Image Lab and quantitation of band intensity was measured with Image J.

### Human Specimen Culturing

Primary human AML specimens were cultured using SFEM II media (Stem Cell Technologies, 09655) supplemented with 10% FBS and 10 nmol/L human cytokines SCF (PEPROTech, 300-07), IL3 (PEPROTech, 200-03), and FLT3 (PEPROTech, 300-19).

### Immunofluorescence microscopy analysis

Leukemia stem cells sorted (using BD FACSAria II) from primary AML specimens were seeded onto fibronectin (RetroNectin - TAKARA BIO INC., T100B) coated glass chamber slides in SFEM II culture medium (described above). Cells were cultured for 1-2 hours at 37 degree Celsius and then fixed in 4% paraformaldehyde for 20 minutes at 4°C, permeabilized with 0.1% Triton X-100 (catalog T9284, Sigma-Aldrich) for 10 minutes at room temperature, and blocked with 5% Normal Donkey Serum (Sigma, 566460) in PBS for one hour. The slides were stained with primary antibodies in 5% normal Donkey serum at 4°C overnight: Rabbit anti-IRF2BP2 (Proteintech, 18847-1-AP). The cells were washed with PBS twice and then labeled with secondary antibodies, Donkey anti-rabbit Alexa Fluor 488 (Invitrogen, catalog A21206) at 1:250 v/v concentration for 2 hours at room temperature. Cells were washed with PBS and then slides were mounted using Prolong Gold Antifade mounting media (Thermo Fisher Scientific, P36935) containing DAPI. The stained cells were acquired using a LSM 780 confocal microscope system (Carl Zeiss) equipped with an inverted microscope (Observer Z1, Zeiss) using a Plan Apochromat ×63 1.4 NA oil immersion lens. Images were analyzed and processed using Image J and Adobe Photoshop v7.

### Colony-forming Assays

Primary AML specimens were plated in human methylcellulose (R&D systems, HSC003) at cell concentrations indicated in respective figures. Samples were treated with DMSO, ven/aza or Triacsin C (Tocris Bioscience, Cat. 76896-80-5) for 16 hours, washed of drug, and added into methylcellulose cultures. Colonies were counted at 12 days after initial plating. <u>Human CD34+ hematopoietic stem and progenitor cells</u> were isolated via FACS using bone marrow harvested from femoral head sample donated from healthy patients. CD34+ cells were cultured in SFEM II base (Stem Cell Technologies, 09655) with 25ng/mL of TPO and SCF, 50ng/mL of Flt3, 10ng/mL of IL3, 0.5μM of SR1, and 1μM of UM729. Cells were treated with DMSO, ven/aza or Triacsin C for 16 hours, washed of drug, and added into methylcellulose (R&D systems, HSC003). Colonies were scored at 12 days after initial plating.

### PDX engraftment

Leukemia stem cell function was assessed by measuring the engraftment of primary AML specimens treated with an ACSL1 inhibitor and transplanted into NSG-S mice. One day prior to transplant, freshly thawed primary AML cells were treated in culture overnight with indicated agents in media with cytokines as listed above. NSG-S mice were conditioned with 25 mg/kg busulfan via i.p. injection. Overnight-treated primary AML cells were washed and resuspended in PBS supplemented with 2% FBS. Anti-human CD3 antibody (BioXCell) was added at a final concentration of 1μg/10^6^ cells to prevent potential graft-versus-host disease. Each mouse recipient received 0.1mL containing 2×10^6^ cells via tail vein injection; there were 8 to 12 mice per experiment group. Mice engrafted with primary AML cells were sacrificed after 4 to 8 weeks. Engraftment was measured by flow cytometry for human CD45+ cells (BD no. 571875, dilution 1:100). All animal studies were done at the University of Colorado under Institutional Animal Care and Use Committee-approved protocol no. 308. The University of Colorado is accredited by the Association for Assessment and Accreditation of Laboratory Animal Care, abides by the Public Health Service Animal Assurance of Compliance, and is licensed by the United States Department of Agriculture.

### Primary AML scRNA-seq Analysis

Differential expression analysis was performed using the EdgeR (v4.2.0) package in R (v.4.4.0) on single-cell data categorized by venetoclax response (sensitive vs. resistant). The dataset used was previously generated and published in Sheth et al., 2024. The full Seurat object was subset to include HSCs & MPPs, Early promyelocytes, Erythro-myeloid progenitors, Late promyelocytes, and Classical Monocytes clusters and only samples with a known response to venetoclax. A pseudobulk object was generated from the Seurat object using the Seurat2PB function.

Samples were annotated based on venetoclax response. Low-quality samples with less than 50,000 unique molecular identifiers (UMIs) were filtered out, and clusters with less than two response levels were excluded from the analysis. A design matrix was constructed for each cluster, modeling venetoclax response as a factor. Genes were then filtered to retain those with a minimum of 5 counts and a minimum total count of 10 across samples. Normalization was carried out using the TMM method, followed by dispersion estimation. The quasi-likelihood F-test from EdgeR’s QL v4 pipeline was used to assess differential expression, with the qlmQLFit function applied to each cluster. The final list of differentially expressed genes was generated from the results of the F-test.

The 1090 significantly differentially expressed genes that overlapped with the IRF2BP2 transcriptional targets were annotated into transcriptional programs based on the results of GO enrichment analysis for biological processes. This was performed using enrichGO from the clusterProfiler R package (v4.12.6) with a p-value cutoff of 0.05 and a q-value cutoff of 0.2. GO pathways were manually filtered into metabolism, differentiation, and inflammation functional programs based on keyword matching in the GO term descriptions. Genes that mapped to multiple programs were assigned to the most biologically relevant program.

### Metabolomics

#### Sample preparation

Primary AML cells were stained and sorted for ROS-low LSC fraction (described above), and 200,000 cells were pelleted per replicate. Quadruplicates for each primary AML were submitted for metabolite extraction and mass spectroscopy analysis. When Triacsin C was used, ROS-low LSCs were treated with 2.5μM Triacsin C for 16 hours, washed of drug and pelleted for metabolite extraction as indicated.

#### Extraction of metabolites from AML cells were as follows

Variable amounts of cold MeOH:ACN:H2O (5:3:2, v:v:v) were added to each cell pellet so that the final extraction concentration of each sample was 2e6 cells/mL. Samples were then vortexed at 4 °C for 30 minutes. Following vortexing, samples were centrifuged at 12700 RPM for 10 minutes at 4 °C and supernatant was transferred to a new autosampler vial for analysis. A portion of extract from each sample was also combined to create a technical mixture, injected throughout the run for quality control.

#### High-throughput Metabolomics Analysis

Analyses were performed as previously published via a modified gradient optimized for the high-throughput analysis of metabolomics.1,2,3 Briefly, the analytical platform employs a Vanquish UHPLC system (Thermo Fisher Scientific, San Jose, CA, USA) coupled online to a Q Exactive mass spectrometer (Thermo Fisher Scientific, San Jose, CA, USA). Metabolomics extracts were resolved over an ACQUITY UPLC BEH C18 column (2.1 × 100 mm, 1.7 µm particle size) held at 45 °C (Waters, MA, USA). For positive mode mobile phase (A) 0.1% FA in water and mobile phase (B) 0.1% FA in ACN was used. For negative mode mobile phase (A) 10mM Ammonium Acetate in Water and mobile phase (B) 10mM Ammonium Acetate in 50:50 ACN:MeOH was used. For negative and positive mode analysis the chromatographic the gradient was as follows: 0.45 mL/min flowrate for the entire run, 0% B at 0 min, 0% B at 0.5 min, 100% B at 1.1 min, 100% B at 2.75min, 0% B at 3min, 0% B at 5 min. For positive ion mode the Exploris 120 mass spectrometer (Thermo Fisher) scanned in Full MS mode from 65 to 975 m/z at 120,000 resolution, with 3.5 kV spray voltage, 50 sheath gas, 10 auxiliary gas. For negative ion mode the Exploris 120 mass spectrometer (Thermo Fisher) scanned in Full MS mode from 65 to 975 m/z at 120,000 resolution, with 3.4 kV spray voltage, 50 sheath gas, 10 auxiliary gas. Calibration was performed prior to analysis using the PierceTM Positive and Negative Ion Calibration Solutions (Thermo Fisher Scientific).

Data analysis: Acquired data was converted from raw to mzXML file format using Mass Matrix (Cleveland, OH, USA). Analysis was done using MAVEN, an open-source software program for metabolomics analysis. Samples were analyzed in randomized order with a technical mixture injected interspersed throughout the run to qualify instrument performance.

### Seahorse Assay

1 million primary AML ROS-low LSCs were treated with 2.5μM Triacsin C and DMSO in SFEM-II, 10ng/μL SCF, 10ng/μL Flt3, 10ng/μL IL3 for 16 hours. Cells were harvested and resuspended in limited seahorse media with 2mM Glucose and 0.5mM Carnitine. 200K cells were plated per well in quadruplicate of a XFe96 plate coated with CellTak (Corning, 354240). Right before running the plate on Seahorse XFe 96 Extracellular Flux Analyzer, cells were supplemented with BSA conjugated palmitate (according to the manufacturer’s instructions - Agilent Technologies-Seahorse XF Palmitate Oxidation Stress Test Kit-103693-100). Oxygen consumption rate (OCR) was measured at basal and after injection of limited media (or 4uM Etomoxir), 5μg/mL oligomycin, 2μM FCCP and 5μM Antimycin A plus 5μM Rotenone.

