## Supplemental Figures S1-5 for "Therapy resistance in AML is mediated by cytoplasmic sequestration of the transcriptional repressor IRF2BP2"

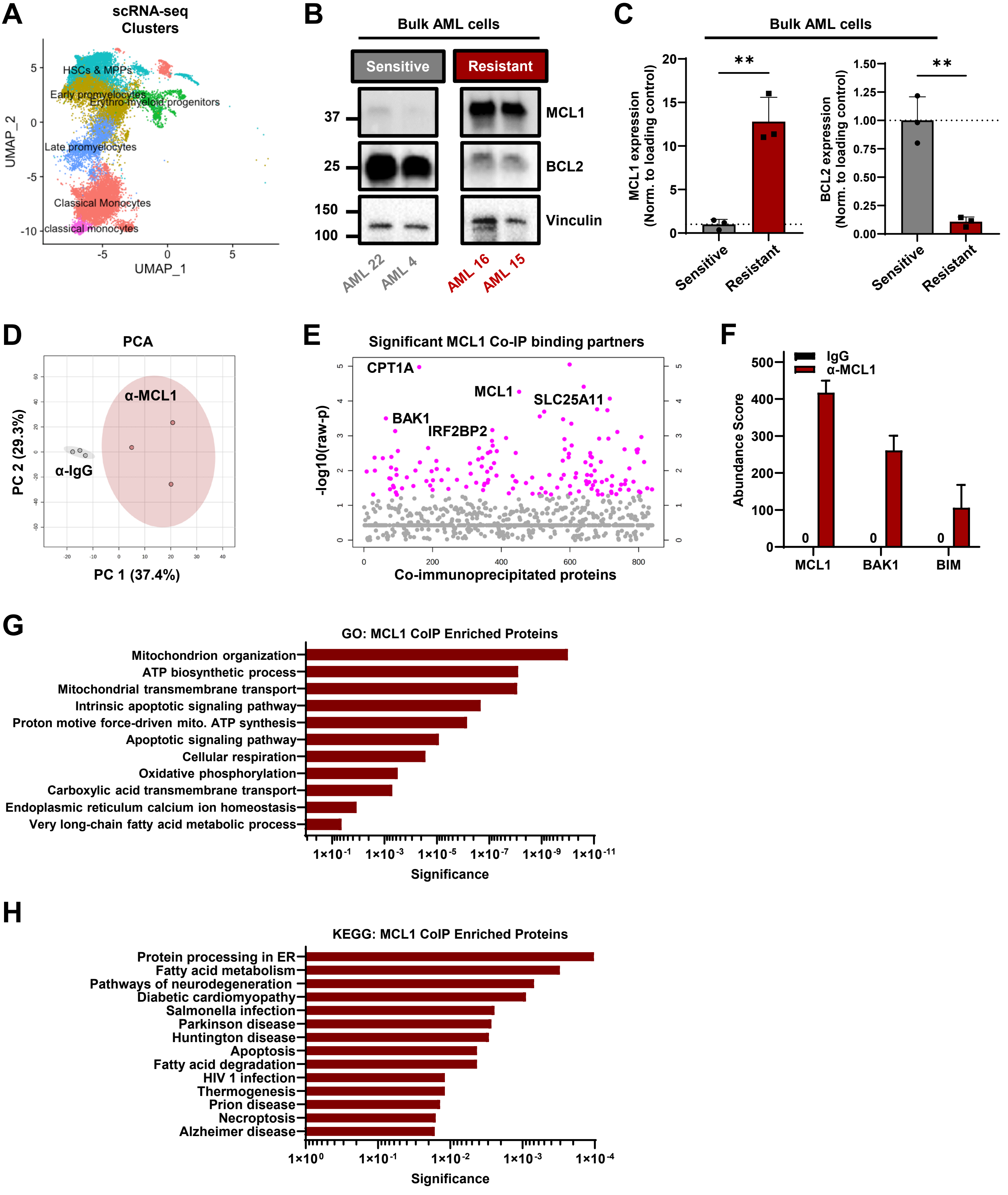


**Supplemental Figure 1**

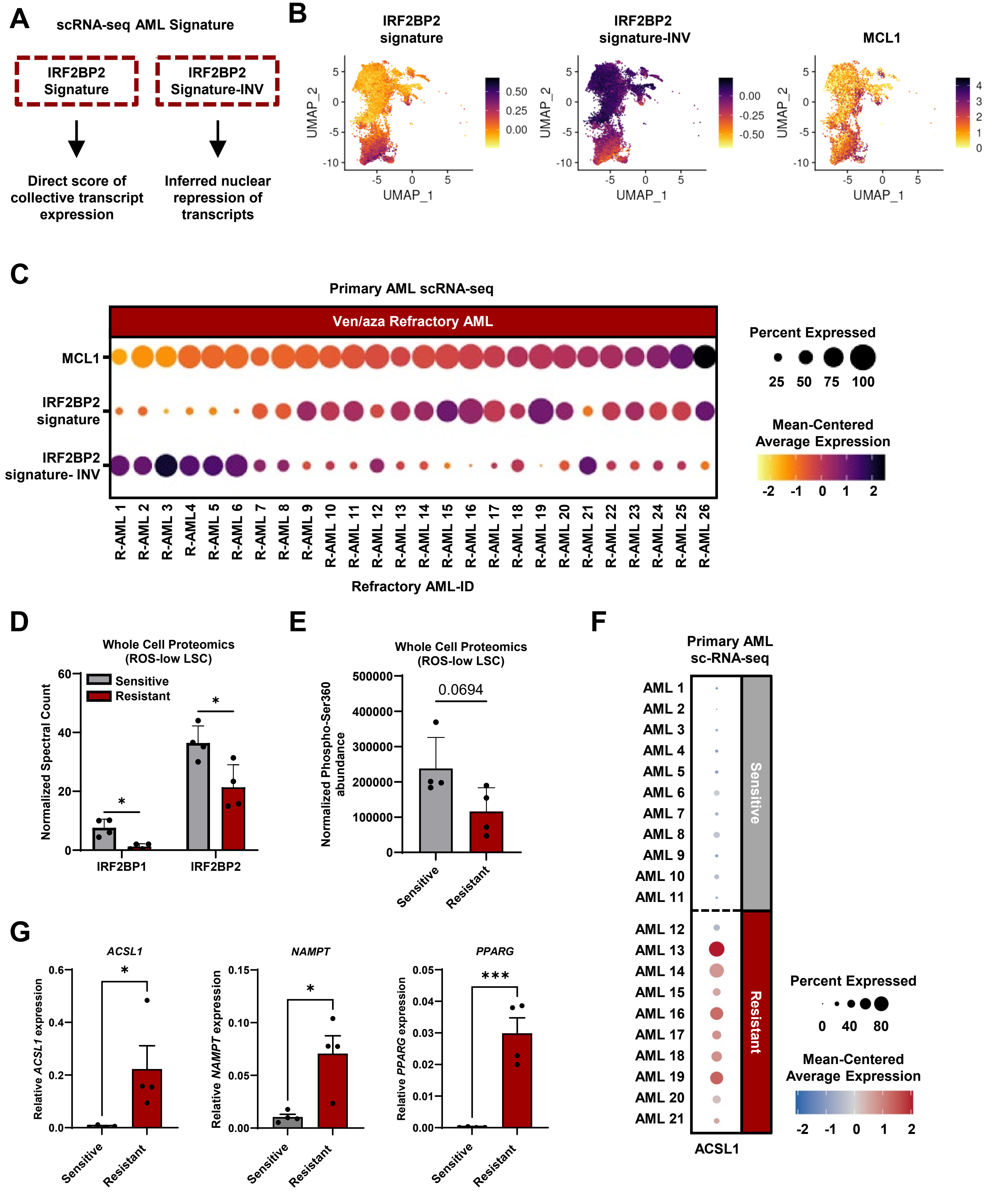


**Supplemental Figure 2**

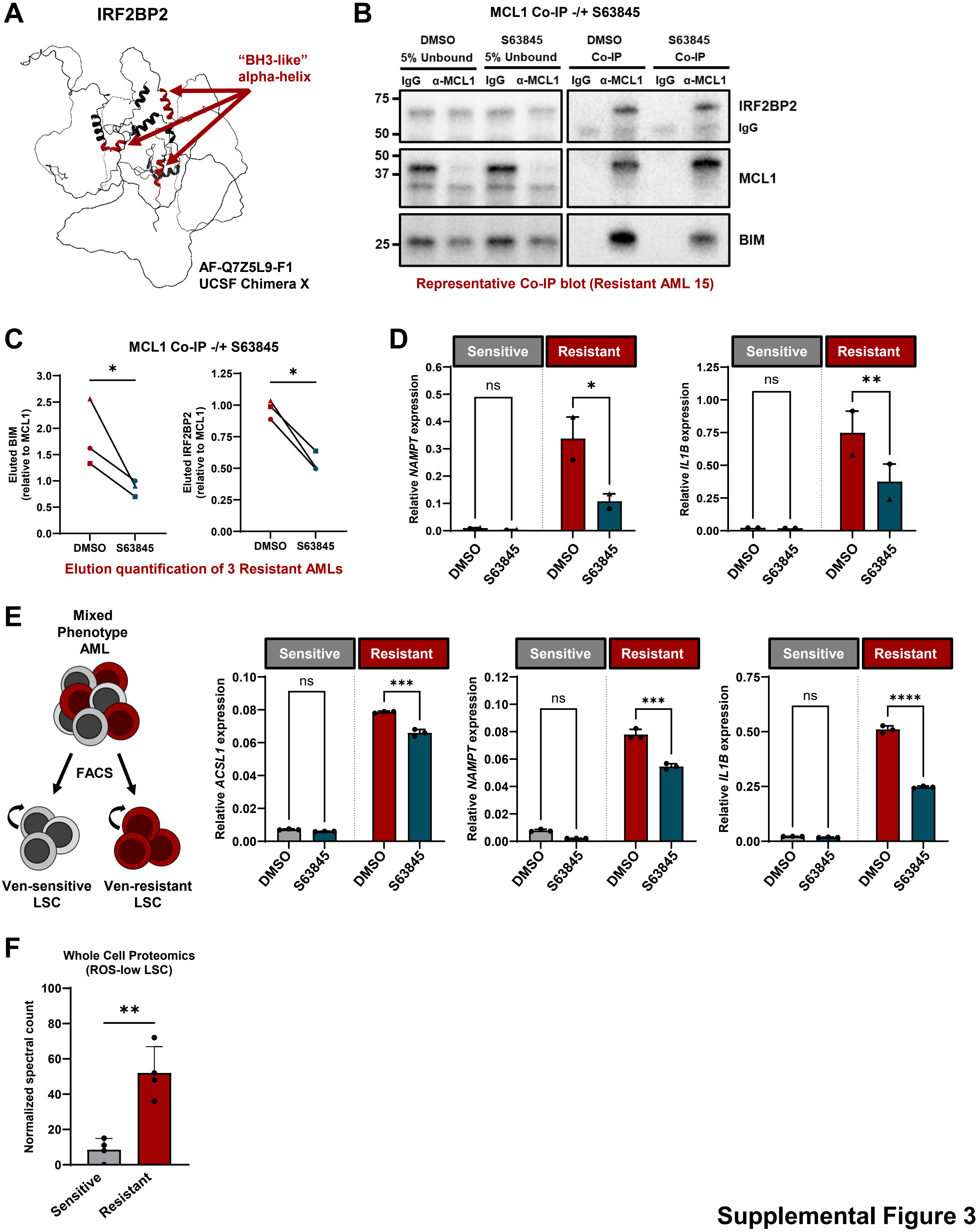


Supplemental Figure 3

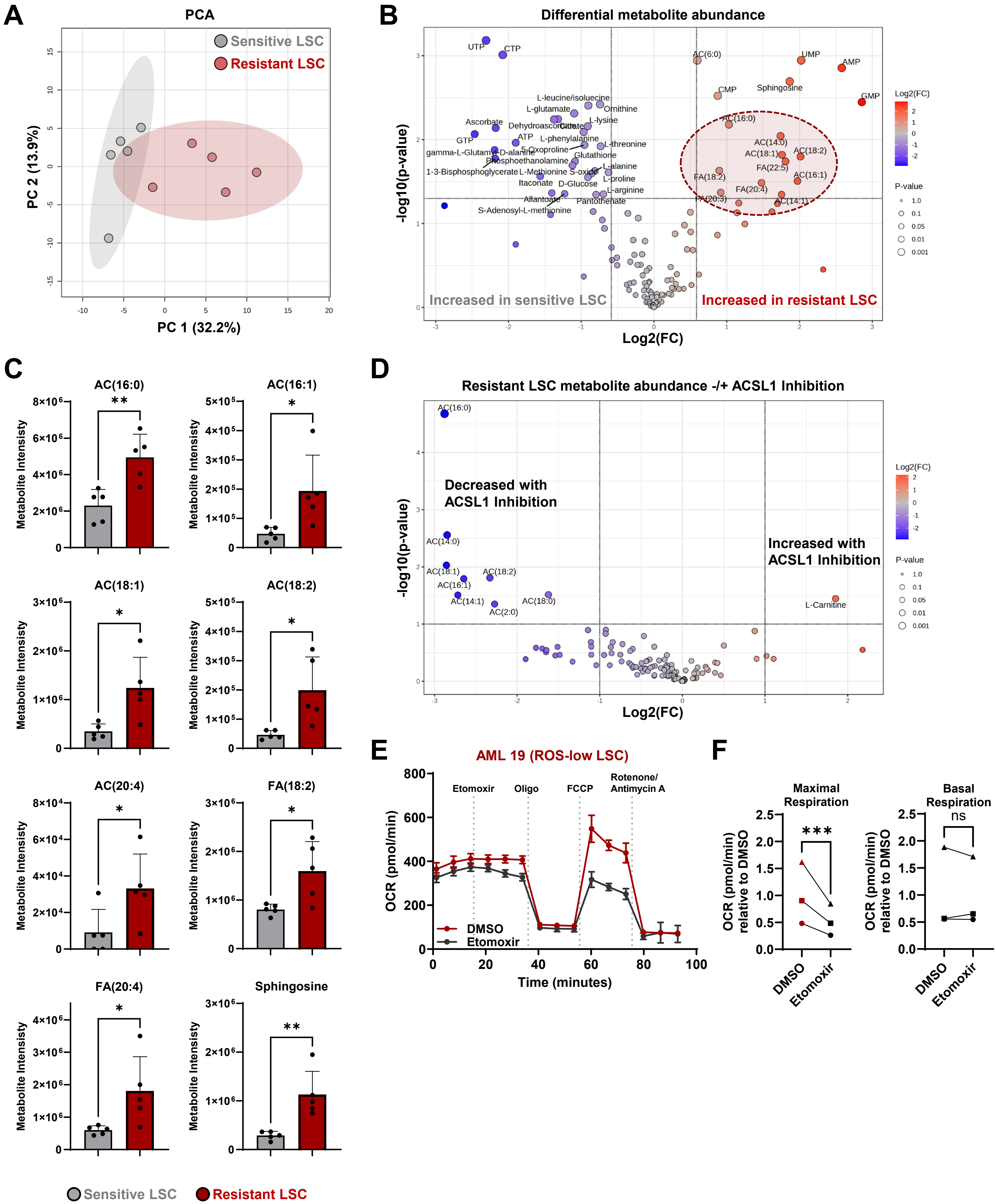


### Supplemental Figure 4

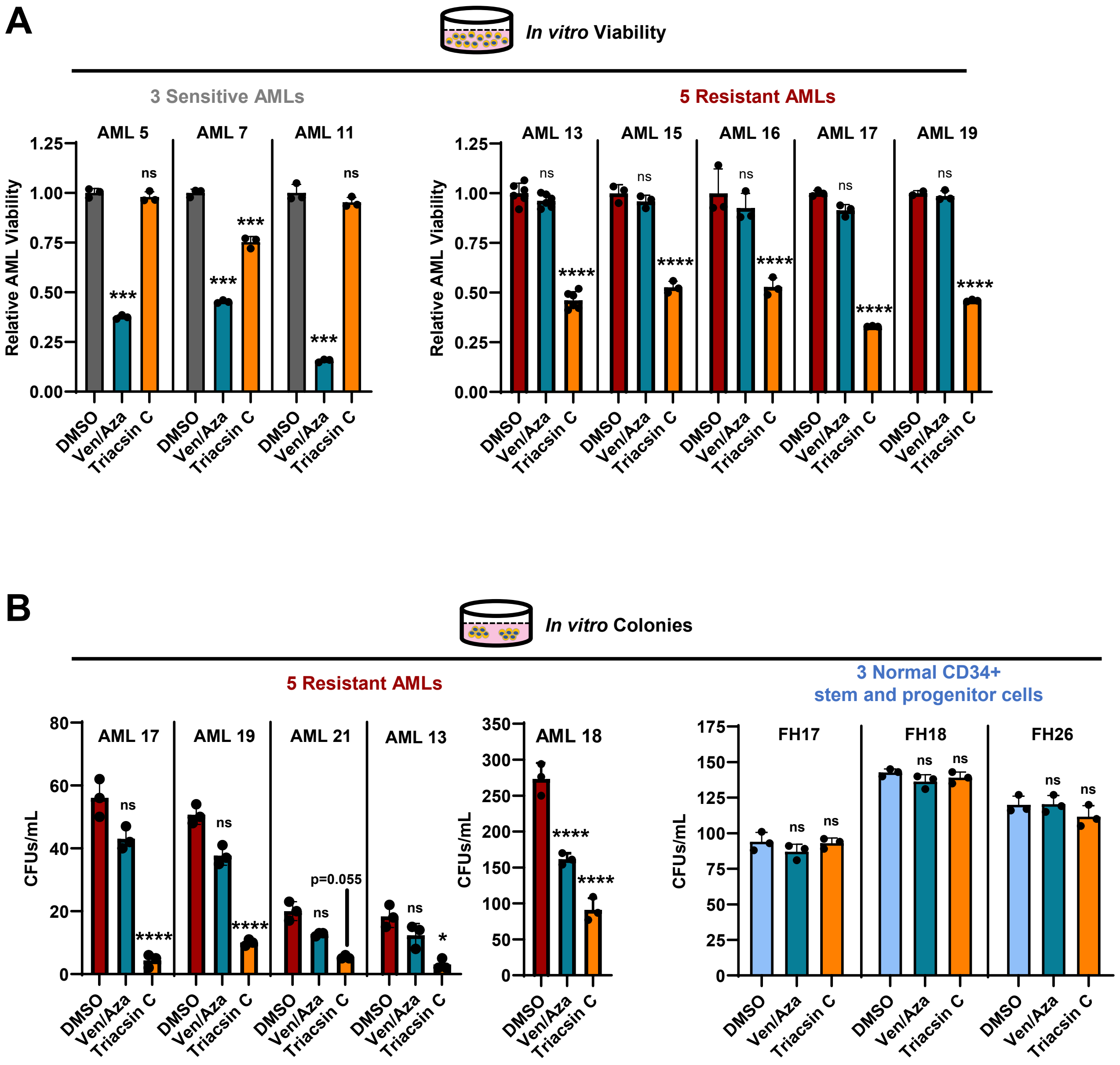


Supplemental Figure 5
