## Supplemental Figure Legends for "Therapy resistance in AML is mediated by cytoplasmic sequestration of the transcriptional repressor IRF2BP2"

**Supplemental Figure 1. (A)** Projection of all primary AML samples with cluster assignments annotated (excluding normal plasma, B and T cell clusters). **(B)** Western blotting analysis for MCL1 and BCL2 protein expression within representative ven/aza-sensitive (N=2) and -resistant (N=2) primary bulk AML samples. **(C)** Quantification of MCL1 and BCL2 protein expression within ven/aza-sensitive (N=3) and -resistant (N=3) primary bulk AML samples. Data were normalized to vinculin loading control and presented as mean ± SD. Significance was determined using a two-tailed unpaired t test, **p<0.01. **(D)** Principal component analysis between elution products from Isotype IgG and α-MCL1 co-immunoprecipitation in ven/aza-resistant primary AML (N = 3). **(E)** Significantly enriched co-immunoprecipitated proteins within α-MCL1 elution relative to Isotype IgG. Significance was determined using a two-tailed unpaired t test, with p<0.05 cutoff. **(F)** Abundance score for canonical apoptosis binding partners of MCL1, BAK1 and BIM. Data are presented as mean ± SD. **(G)** Gene Ontology pathway analysis of MCL1 co-immunoprecipitated proteins within primary ven/aza-resistant AML (N=3). Depicted are 11 representative biological processes from a total of 147 significant results. Statistical enrichment and significance were determined using hypergeometric testing with a padj<0.05 cutoff. **(H)** KEGG pathway analysis of MCL1 co-immunoprecipitated proteins within primary ven/aza-resistant AML (N=3). Depicted are 14 representative pathways from a total of 16 significant results. Statistical enrichment and significance were determined using hypergeometric testing with a padj<0.05 cutoff.

**Supplemental Figure 2. (A)** Schematic representing primary AML scRNA-seq signature nomenclature. IRF2BP2 signature (107 genes, Ellegast et al., 2022) represents the direct score of the collective transcript expression while IRF2BP2 signature-INV represents inferred nuclear repression of IRF2BP2 target transcripts. **(B)** Projection of all scRNA-seq primary AML samples (from diseased clustering) queried for *MCL1*, IRF2BP2 signature or IRF2BP2 signature-INV. **(C)** Dot plot representing expression and proportion for *MCL1*, IRF2BP2 signature and IRF2BP2 signature-INV within an AML diseased cluster from scRNA-sequencing analysis of primary patient samples clinically refractory to ven/aza therapy (R-AML, N=26). **(D)** IRF2BP1 and IRF2BP2 protein expression from whole cell proteomics analysis comparing ven/aza-sensitive (N=4) and-resistant (N=4) AML ROS-low LSC. Data are presented as mean ± SD. Significance was determined using a two-tailed unpaired t test, *p<0.05. **(E)** Phospho-Serine 360 peptide abundance detected from whole cell proteomics analysis comparing ven/aza-sensitive (N=4) and-resistant (N=4) AML ROS-low LSC. Data are presented as mean ± SD. **(F)** Dot plot representing *ACSL1* expression and proportion within an AML diseased cluster from scRNA-sequencing analysis of respective ven/aza-sensitive (N=11) and -resistant (N=10) primary AML samples. **(G)** Quantitative RT-PCR analysis of *ACSL1*, *PPARG* and *NAMPT* transcript within ven/aza-sensitive (N=4) and -resistant (N=4) AML ROS-low LSC. Data are presented as mean ± SD relative to gene loading control. Significance was determined using a two-tailed unpaired t test, *p<0.05, ***p<0.001.

**Supplemental Figure 3. (A)** 3-D rendering of IRF2BP2 predicted AlphaFold structure within UCSF Chimera X software. Red alpha-helices indicate the putative BH3-like motifs of IRF2BP2. **(B)** Representative western blot for MCL1 coimmunoprecipitation from primary ven/aza-resistant AML subjected to vehicle control or S63845 treatment. Blot was probed for MCL1, IRF2BP2 and BIM. **(C)** Quantification of BIM and IRF2BP2 co-immunoprecipitation relative to MCL1 capture. Significance was determined using a two-tailed ratio paired t test, *p<0.05. **(D)** Relative *NAMPT* and *IL1B* mRNA expression in ven/aza-sensitive (N=2) and -resistant (N=2) ROS-low LSC subjected to vehicle control or S63845 treatment for 4hr. Data are presented as mean ± SD relative to gene loading control. Significance was determined using a two-way ANOVA test, *p<0.05, **p<0.01. **(E)** Relative *ACSL1*, *NAMPT* and *IL1B* mRNA expression in ven/aza-sensitive and -resistant ROS-low LSC isolated from the same primary AML subjected to vehicle control or S63845 treatment for 4hr. Data are presented as mean ± SD relative to gene loading control. Significance was determined using a two-way ANOVA test, *p<0.05, **p<0.01. **(F)** ACSL1 protein expression from whole cell proteomics analysis comparing ven/aza-sensitive (N=4) and-resistant (N=4) primary AML ROS-low LSC. Data are presented as mean ± SD. Significance was determined using a two-tailed unpaired t test, **p<0.01.

**Supplemental Figure 4. (A)** Principal component analysis of differential metabolites between ven/aza-sensitive (N=5) and -resistant (N=5) primary AML ROS-low LSC. **(B)** Volcano plot representing differential metabolite abundance accessed by log2(FC) and -log10(p-value) between ven/aza-sensitive (N=5) and -resistant (N=5) primary AML ROS-low LSC. Encircled are significant increases of long-chain fatty acids and acyl-carnitine species within ven/aza-resistant LSC. **(C)** Long-chain fatty acid, acyl-carnitine, and sphingosine abundance from metabolomic analysis between ven/aza-sensitive (N=5) and -resistant (N=5) primary AML ROS-low LSC. Data are presented as mean ± SD. Significance was determined using a two-tailed unpaired t test, *p<0.05, **p<0.01. **(D)** Volcano plot representing differential metabolite abundance accessed by log2(FC) and -log10(p-value) between ven/aza-resistant (N=3) primary AML ROS-low LSC subjected to vehicle control or Triacsin C treatment for 16 hours. **(E)** Representative oxygen consumption rate (OCR) curve from Seahorse XF Palmitate Oxidation Stress assay comparing respiration of ven/aza-resistant primary AML ROS-low LSC subjected to acute Etomoxir treatment. Technical replicates of four per data point. Data are presented as mean ± SD for 100k cells. Vertical dotted lines indicate injection times used throughout assay. **(F)** Maximal and basal respiration rates calculated from Seahorse XF Palmitate Oxidation Stress assay from ven/aza-resistant (N=3) primary AML ROS-low LSC subjected to acute Etomoxir treatment. Data are presented as OCR relative to DMSO. Significance was determined using a two-tailed ratio paired t test, ***p<0.001.

**Supplemental Figure 5. (A)** *In vitro* viability of ven/aza-sensitive (N=3) and -resistant (N=5) primary AML subjected to ven/aza or Triacsin C treatment for 24 hours. Data are presented as mean ± SD of technical replicates for individual primary AML. Significance was determined using two-way ANOVA test, ***p<0.001, ****p<0.0001. **(B)** *In vitro* colony forming potential of ven/aza-resistant (N=5) primary AML and normal CD34+ hematopoietic stem and progenitor cells (N=3) subjected to ven/aza or Triacsin C treatment for 16 hours. Data are presented as mean ± SD of technical replicates for individual primary AML or normal HSPC. Significance was determined using two-way ANOVA test, *p<0.05, **p<0.01, ***p<0.001, ****p<0.0001.
