## Supplemental Table 1 for "Therapy resistance in AML is mediated by cytoplasmic sequestration of the transcriptional repressor IRF2BP2"

| **Supplemental Table 1. Top MCL1 Co-immunoprecipitated features ranked by abundance score.** | | | | | | | | | |
| --- | --- | --- | --- | --- | --- | --- | --- | --- | --- |
|  | | **α-IgG** | | | **α-MCL1** | | | | **α-MCL1** |
| **#** | **Feature ID** | **AML 15** | **AML 16** | **AML 17** | **AML 15** | **AML 16** | | **AML 17** | **Mean** |
| 1 | IRF2BP2 | 0.00 | 0.00 | 0.00 | 649.18 | | 557.38 | 754.92 | 653.83 |
| 2 | MCL1 | 0.00 | 0.00 | 0.00 | 448.65 | | 417.57 | 383.78 | 416.67 |
| 3 | BAK1 | 0.00 | 0.00 | 0.00 | 306.52 | | 234.78 | 241.30 | 260.87 |
| 4 | IRF2BPL | 0.00 | 0.00 | 0.00 | 216.87 | | 204.82 | 291.57 | 237.75 |
| 5 | GOLGA3 | 0.00 | 0.00 | 0.00 | 128.74 | | 335.93 | 144.01 | 202.89 |
| 6 | TUBB3 | 0.00 | 0.00 | 0.00 | 173.00 | | 287.00 | 61.00 | 173.67 |
| 7 | BCL2L11 | 0.00 | 0.00 | 0.00 | 152.27 | | 129.55 | 36.36 | 106.06 |
| 8 | LMNB1 | 0.00 | 0.00 | 0.00 | 121.21 | | 42.42 | 130.30 | 97.98 |
| 9 | IRF2BP1 | 0.00 | 0.00 | 0.00 | 70.97 | | 90.32 | 103.23 | 88.17 |
| 10 | MYO1E | 0.00 | 0.00 | 0.00 | 53.15 | | 83.07 | 116.54 | 84.25 |
| 11 | TUBB6 | 0.00 | 0.00 | 0.00 | 55.00 | | 116.00 | 32.00 | 67.67 |
| 12 | BBC3 | 0.00 | 0.00 | 0.00 | 66.67 | | 40.48 | 21.43 | 42.86 |
| 13 | DOCK2 | 0.00 | 0.00 | 0.00 | 13.21 | | 71.93 | 30.66 | 38.60 |
| 14 | ATP5MF | 0.00 | 0.00 | 0.00 | 36.36 | | 63.64 | 13.64 | 37.88 |
| 15 | TMEM43 | 0.00 | 0.00 | 0.00 | 47.78 | | 4.44 | 52.22 | 34.81 |
| 16 | RCOR1 | 0.00 | 0.00 | 0.00 | 14.15 | | 32.08 | 51.89 | 32.70 |
| 17 | ATP5ME | 0.00 | 0.00 | 0.00 | 43.75 | | 50.00 | 0.00 | 31.25 |
| 18 | RAB44 | 0.00 | 0.00 | 0.00 | 38.29 | | 18.02 | 36.04 | 30.78 |
| 19 | RHOG | 0.00 | 0.00 | 0.00 | 38.10 | | 26.19 | 26.19 | 30.16 |
| 20 | HSD17B11 | 0.00 | 0.00 | 0.00 | 24.24 | | 36.36 | 21.21 | 27.27 |
| 21 | BTBD2 | 0.00 | 0.00 | 0.00 | 20.54 | | 10.71 | 49.11 | 26.79 |
| 22 | DYNLL2 | 0.00 | 0.00 | 0.00 | 20.00 | | 60.00 | 0.00 | 26.67 |
| 23 | KDM1A | 0.00 | 0.00 | 0.00 | 8.06 | | 20.43 | 49.46 | 25.99 |
| 24 | SFXN3 | 0.00 | 0.00 | 0.00 | 29.17 | | 33.33 | 13.89 | 25.46 |
| 25 | ERLIN2 | 0.00 | 0.00 | 0.00 | 35.53 | | 11.84 | 25.00 | 24.12 |
| 26 | CHCHD3 | 0.00 | 0.00 | 0.00 | 50.00 | | 7.69 | 13.46 | 23.72 |
| 27 | BTBD1 | 0.00 | 0.00 | 0.00 | 16.04 | | 9.43 | 41.51 | 22.33 |
| 28 | IMMT | 0.00 | 0.00 | 0.00 | 23.21 | | 14.88 | 28.57 | 22.22 |
| 29 | BMF | 0.00 | 0.00 | 0.00 | 26.19 | | 23.81 | 16.67 | 22.22 |
| 30 | CRACR2A | 0.00 | 0.00 | 0.00 | 24.70 | | 15.06 | 26.51 | 22.09 |
| 31 | ERLIN1 | 0.00 | 0.00 | 0.00 | 28.21 | | 15.38 | 19.23 | 20.94 |
| 32 | RIPK3 | 0.00 | 0.00 | 0.00 | 20.18 | | 20.18 | 21.93 | 20.76 |
| 33 | TMEM33 | 0.00 | 0.00 | 0.00 | 26.79 | | 19.64 | 14.29 | 20.24 |
| 34 | HSPD1 | 0.00 | 0.00 | 0.00 | 3.28 | | 31.15 | 26.23 | 20.22 |
| 35 | SKP1 | 0.00 | 0.00 | 0.00 | 7.89 | | 31.58 | 21.05 | 20.18 |
| 36 | CPT1A | 0.00 | 0.00 | 0.00 | 20.45 | | 18.75 | 18.75 | 19.32 |
| 37 | TOMM22 | 0.00 | 0.00 | 0.00 | 31.25 | | 15.63 | 9.38 | 18.75 |
| 38 | DPYSL2 | 0.00 | 0.00 | 0.00 | 19.35 | | 20.97 | 12.10 | 17.47 |
| 39 | HDAC1 | 0.00 | 0.00 | 0.00 | 3.64 | | 17.27 | 30.00 | 16.97 |
| 40 | RAB32 | 0.00 | 0.00 | 0.00 | 16.00 | | 18.00 | 14.00 | 16.00 |
| 41 | LMNA | 0.00 | 0.00 | 0.00 | 21.62 | | 10.14 | 15.54 | 15.77 |
| 42 | FBXO11 | 0.00 | 0.00 | 0.00 | 7.21 | | 10.10 | 28.37 | 15.22 |
| 43 | RPS19 | 0.00 | 0.00 | 0.00 | 12.50 | | 9.38 | 18.75 | 13.54 |
| 44 | KIF5B | 0.00 | 0.00 | 0.00 | 9.55 | | 11.82 | 18.64 | 13.33 |
| 45 | RNASE3 | 0.00 | 0.00 | 0.00 | 8.33 | | 11.11 | 19.44 | 12.96 |
| 46 | CYC1 | 0.00 | 0.00 | 0.00 | 11.43 | | 8.57 | 18.57 | 12.86 |
| 47 | BCL2 | 0.00 | 0.00 | 0.00 | 17.31 | | 9.62 | 11.54 | 12.82 |
| 48 | PHB | 0.00 | 0.00 | 0.00 | 21.67 | | 6.67 | 10.00 | 12.78 |
| 49 | ENO3 | 0.00 | 0.00 | 3.19 | 22.34 | | 0.00 | 15.96 | 12.77 |
| 50 | HDAC2 | 0.00 | 0.00 | 0.00 | 0.00 | | 11.82 | 26.36 | 12.73 |
