## Supplemental Table 2 for "Therapy resistance in AML is mediated by cytoplasmic sequestration of the transcriptional repressor IRF2BP2"

| **Supplemental Table 2. Characteristics of Patient Specimens** | | | | | |  |
| --- | --- | --- | --- | --- | --- | --- |
| **AML ID** | **Diagnosis** | **Age** | **Sex** | **Karyotyping** | **Mutations** | **Ven/aza Sensitivity** |
| AML 1 | De Novo | 73 | F | 45,XX,add(3)(q27),-7, add(8)(q24),t(9;22)(q34;q11.2)[4]/54,sl,+3,-add(3)(q27),+7,+8,+10,+12,+13,+15,+20,+der(22)t(9;22)(q34;q11.2)[16] |  | Sensitive |
| AML 2 | De Novo | 52 | M | 45,XY,-7[3]/46,sl,+r(7)(p11q21)[11]/46,sdl1,der(5)t(1;5)(q31;p14)[5]/46,XY[1] | ASXL1 c.4120_4121insT;  DNMT3A c.885delG;  Notch1 c.4746_4747insTGGGGA;  NRAS c. 182A>G | Sensitive |
| AML 3 |  |  |  | clone 1: +8q, +3q, t(9;22); clone 2: clone 1 abnormalities AND +3, +10, +12 +13, +15 +20; clone 3: clone 2 abnormalities AND der (22); by FISH -7 | FLT3-ITD+, WT for NPM1, CEBPA, IDH1 and IDH2 | Sensitive |
| AML 4 | De Novo |  |  |  | TET2 c.T5162G; p.L1721W (99.2%), NRAS c.G37C; p.G13R (30.2%), ASXL1 c.T2444C; p.L815P (99.8%), GATA2 c.T962A; p.L321H (52.0%), BCORL1 c.T331C; p.F111L(100%), BCORL1 c.C2912G; p.A971G (99.8%) | Sensitive |
| AML 5 | Relapse | 47 | M | 46,XY,del(7)(q21)[8]/46,sl,del(5)(q31q35),add(12)(p13)[7]/46,sl,add(12)(p13),del(17)(q21)[3]/46,XY,del(9)(q22q32)[2] | IDH1 R132; CKIT D816V | Sensitive |
| AML 6 | De Novo | 80 | F | 47,XX,+8[15]/47,XX,+11[1]/44-45,XX,add(5)(p12),dic(5;12)(q10;q10),t(11;14)(q13;q32),dic(17;18) (q10;q10), add(20)(q11.2) [cp8]/44,XX,dic(5;12)(q10;q10), der(8)(11qter- > 11q13::14q32- > 14q10::8p11.2- > 8qter), der(11)t(11;14)(q13;q32), dic(17;18)(q10;q10)[2]/46,XX[5] | CBL M400R, SRSF2 P95H, TET2 Q916X and S1290X | Sensitive |
| AML 7 | Relapse | 56 | M | 46,XY[20] normal | NRAS.G13C 13%, GATA2.L375I 42%, KRAS.G12V 31%, CEBPA.P49Rfs*111 42% | Sensitive |
| AML 8 | De Novo |  | F | 46,XX,t(6;11)(q27;q23)[18]/92,slx2[2] | KRAS.G12C 16%,  PTPN11.E76L 7% | Sensitive |
| AML 9 | Relapse | 36 |  | normal | TET2.I1762V, ASXL1.L815P, CUX1.A448T,  BCORL1.F111L, TP53.P33R | Sensitive |
| AML 10 | Relapse | 49 | F | 46, XX normal | FLT3 ITD+, FLT3 D835 | Sensitive |
| AML 11 | De Novo | 50 | M | 46,XY,add(1)(p11),del(5)(q15q33),del(7)(q22q36),der(11)t(1;11)(p31;p12-14)[20] | FLT3 ITD+ (38%), BCOR F1023L (15%), NOTHC1 S2542N (12%) | Sensitive |
| AML 12 | Relapse | 79 | M | 46,XY,t(6;9) (p21;q34) | FLT3 ITD+ and FLT3 TKD mutation (D835); IDH2 R140Q; | Resistant |
| AML 13 | De Novo | 74 | F | 46,XX[20] normal | FLT3 TKD (D835), NPM1+, TET2 c.4311_4318GAAAAAC, 41%, TET2.I1762V 24%, PTPN11.A72T 43%, ASXL1.L815P 100%, BCCORL1.F111L 100%, DNMT3AA.G354C 43%, CUX1.A448T 100%, NOTCH1.V2285I 41%, CBLC.P389S 38%. | Resistant |
| AML 14 | De Novo | 52 | M | 45,XY,-7[3]/46,sl,+r(7)(p11q21) [11]/46,sdl1,der(5)t(1;5)(q31;p14)[5]/46,XY[1] | ASXL1 c.4120_4121insT;  DNMT3A c.885delG;  Notch1 c.4746_4747insTGGGGA; NRAS c. 182A>G | Resistant |
| AML 15 | Relapse | 65 | F | 46,XX,add(14)(q22)[4], 46,XX[16] | FLT3+, NPM1+, IDH1+ | Resistant |
| AML 16 | De Novo | 63 | M | 46,XY[19] normal | FLT3 ITD+, NPM1+ | Resistant |
| AML 17 | De Novo | 60 | F | 46,XX,t(9;11)(p21;q23)[13]/47,sl,+21[4]/47,sl,+8[3] | KRAS.G13D 23%, KRAS.A146T 6%, PTPN11.T52A 47% | Resistant |
| AML 18 | CMML progressed to AML | 69 | M | 46,XX[16] normal | NPM1.W288CFs*12 37%, ASXL1.G646Wfs*12 36%, NRAS.G12D 46%, STAG2.G46Rfs*41 11%, EZH2.Q533L 50%, EZH2.L149Q 49%, TET2.N1610Ifs*6 50%, TET2.Q1699* 50% | Resistant |
| AML 19 | De Novo | 63 | M | 46,XY[18] | DNMT3A.G550R 40%, DNMT3A.splice site variant 18%, NPM1.W288Cfs*12 22%, ETV6.R103_Y104delinsH 15%, KRAS.G13D 14%, GATA2.R343G 35% | Resistant |
| AML 20 | De Novo |  | F | 46,XX,t(6;11)(q27;q23)[18]/92,slx2[2] | KRAS.G12C 16%, PTPN11.E76L 7% | Resistant |
| AML 21 | De Novo | 77 | F | 46,XX[20] normal | NPM1: W288fs 45%, RUNX1: S469P 6%, TET2: V1232del 48% | Resistant |
| AML 22 | De Novo |  |  | t(16;21) |  | Sensitive |
| AML 23 | De Novo | 50 | M | 45,X,-Y; t(9;11)(p22;q23) | FLT3 ITD+ | Resistant |
| AML 24 | Relapse | 69 | M | 46,XY,t(5;12)(q32;p13)[20] | FLT3-ITD (91%), NPM1 (44%), WT1 (22%), DNMT3A (42%) | Sensitive |
